# Intrinsic Heterogeneity of Primary Cilia Revealed Through Spatial Proteomics

**DOI:** 10.1101/2024.10.20.619273

**Authors:** Jan N. Hansen, Huangqingbo Sun, Konstantin Kahnert, Eini Westenius, Alexandra Johannesson, Carmela Villegas, Trang Le, Kalliopi Tzavlaki, Casper Winsnes, Emmie Pohjanen, Anna Mäkiniemi, Jenny Fall, Frederic Ballllosera Navarro, Anna Bäckström, Cecilia Lindskog, Fredric Johansson, Kalle von Feilitzen, Angelica M. Delgado-Vega, Anna Martinez Casals, Diana Mahdessian, Mathias Uhlén, Shu-Hsien Sheu, Anna Lindstrand, Ulrika Axelsson, Emma Lundberg

**Affiliations:** Science for Life Laboratory, School of Engineering Sciences in Chemistry, Biotechnology and Health, KTH - Royal Institute of Technology, Stockholm, Sweden; Department of Bioengineering, Stanford University, Stanford, CA, USA; Department of Molecular Medicine and Surgery, Karolinska Institutet, Stockholm, Sweden; Department of Clinical Genetics and Genomics, Karolinska University Hospital, Stockholm, Sweden; Chan Zuckerberg Imaging Institute, Redwood City, CA, USA; Department of Immunology, Genetics and Pathology, Cancer Precision Medicine Research Unit, Uppsala University, Uppsala, Sweden; Department of Neuroscience, Karolinska Institutet, Stockholm, Sweden; Department of Pathology, Stanford University, Stanford, CA, USA; Chan Zuckerberg Biohub, San Francisco, San Francisco, CA, USA

**Keywords:** Primary Cilia, Cilia, Cellular Heterogeneity, Spatial Proteomics, Immunofluorescence Microscopy, Ciliopathies, 3D Images

## Abstract

Primary cilia are a critical organelle found on most human cells, and their dysfunction is linked to hereditary ciliopathies with a wide phenotypic spectrum. Despite their significance, the specific roles of cilia in different cell types remain poorly understood due to limitations in analyzing ciliary protein composition. We employed antibody-based spatial proteomics to expand the Human Protein Atlas to primary cilia. Our analysis identified the subciliary locations of 715 proteins across three cell lines, examining 128,156 individual cilia. We found that 69% of the ciliary proteome is cell-type specific, and 78% exhibited single-cilia heterogeneity. Our findings portray cilia as sensors tuning their proteome to effectively sense the environment and compute cellular responses. We identified 91 novel cilia proteins and found a genetic candidate variant in *CREB3* in one clinical case with features overlapping ciliopathy phenotypes. This open, spatial cilia atlas advances research on cilia and ciliopathies.

## Introduction

Cilia are small, hair-like cellular appendages that exist on almost any vertebrate cell. They are classified into motile cilia and immotile, primary cilia. While motile cilia move entire cells or fluid flow along epithelial surfaces, primary cilia are immotile and fulfill an antenna function ^1^. Most of what we know about the function of primary cilia stems from hereditary diseases that are caused by primary cilia dysfunction, so-called ciliopathies ^2^. Ciliopathy patients can present with a broad, heterogeneous spectrum of overlapping phenotypes, which include polydactyly, cognitive impairments, obesity, kidney defects, retinal degeneration and blindness, skeletal dysplasia, and endocrine abnormalities, underlining that primary cilia fulfill pleiotropic and tissue-specific functions ^2^. However, the molecular mechanisms underlying primary cilia function are largely unknown.

Functionally, the primary cilium represents an organelle optimized for signal transduction: they sense extracellular cues via receptors, locally transduce the signal, and relay it to the cell body ^3^. The primary cilium reportedly participates in several signaling pathways: individual components of Hedgehog signaling, G-protein-coupled receptor (GPCR) signaling, receptor-tyrosine-kinase (RTK) signaling as well as transforming growth factor β (TGF-β) signaling, bone morphogenetic protein (BMP) signaling, and Wnt signaling have been observed to localize to the primary cilium^1^. However, while the molecular sequence of events has been mostly understood only for Hedgehog signaling ^1^, knowledge about how the other components engage mechanistically in signaling pathways and whether and how the cilium can distinguish between signals of different receptors targeting the same signaling mechanism (e.g., GPCRs) remains elusive ^4^.

The architecture of the cilium and the process of its construction, called ciliogenesis, have been extensively studied. Two ciliogenesis pathways have been described, the external and the internal pathway, which are initiated by the mother centriole docking to the plasma membrane or a vesicular membrane, respectively ^5^. Next, the cilium emerges by extension of the axoneme from the mother centriole. The axoneme is the central skeleton structure of the cilium consisting of nine microtubule doublets nucleated at the centriole. Beyond providing the cilium with a skeleton and thereby determining its length, the axoneme serves as a template for protein trafficking in the cilium, which is driven by the intraflagellar transport (IFT) machinery ^1^. Early during ciliogenesis, several protein structures are built around the mother centriole; The centriole becomes the “basal body” ^5^. Transition fibers are formed to mount the basal body to the plasma membrane ^6–9^. At the interface of cilium and cell body, the “transition zone” serves as a gate limiting the exchange of lipids and proteins between the cilium and the cell body, which macro-structurally appear as a continuum ^10^. The transition zone consists of several protein structures, including Y-shaped protein complexes connecting the plasma membrane and ciliary axoneme ^11^ or the distal appendage matrix sealing the space between transition fibers ^12^. Yet, the composition and mechanisms of the gate between the cilium and cell body remain incompletely understood.

Nonetheless, it has been shown that the protein and lipid repertoire of the primary cilium is distinct from the cell body, which is achieved by controlled entry and exit of proteins ^13,14^. This tightly controlled shuttling of proteins into and out of the primary cilium fulfills multiple functions: (1) Enabling signal transduction, e.g., during activation of Hedgehog signaling a competitive spatial exchange of proteins triggers the signaling cascade ^15^. (2) Relaying information to the cell body to regulate cellular responses, e.g., in Hedgehog signaling, transcription factors are modified in the primary cilium, shuttle to the nucleus and instruct a specific gene expression program ^15^. (3) Tuning the sensitivity of the ciliary antenna function, e.g., activated GPCRs in the primary cilium are removed from the ciliary membrane and transported out of the cilium ^16,17^, and how ciliary length determines its sensitivity in mechanosensation ^18^. (4) Assembling and disassembling the primary cilium ^5^. In conclusion, understanding the ciliary protein repertoire and the underlying spatiotemporal dynamics appears key to understanding how the cilium is influenced by and shaping cell states ^1^.

A major limitation for primary cilia research is imposed by the size of the primary cilium. Its volume is in the range of 0.35 fl, which is approximately 5,000-times smaller than the cell body ^19^, thus requiring highly sensitive measurement technologies. In addition, extracting and purifying intact cilia is difficult, if not impossible, because the ciliary membrane is a continuum of the plasma membrane ^5^. Thus, the ciliary proteome has mainly been characterized using proximity-labeling-based spatial proteomics techniques, such as APEX-, BioID, or Turbo-ID-based proximity labeling ^20–25^. While these methods offer time-efficient, deep, and quantitative studies of the ciliary protein composition, proximity-labeling MS methods face certain limitations: (1) they do not provide single cell resolution and, thus, results represent a mix of cilia in multiple signaling states, and the accuracy of the method is largely influenced by the ciliation rate of the studied cells; (2) technologically, they can only reveal ciliary protein enrichment compared to the cell body; (3) they are not functional in all cell types due to endogenous enzymatic activity; (4) they rely on fusing an enzyme, such as APEX or TurboID, to a ciliary GPCR or ciliary protein fragment, which may bias signaling in the primary cilium; (5) their sensitivity is limited in cell types with high endogenous peroxidase activity ^26^; and (6) they are not applicable to study human tissues.

Here, we tackle these challenges and the concomitant gaps in our understanding of the ciliary proteome and shed light on the physiological function of individual primary cilia on human cells by applying large-scale antibody- and imaging-based spatial proteomics. We establish a new resource for the cilia field: a section in the public Human Protein Atlas (HPA) that provides high-resolution confocal 3D images and subciliary annotations for 715 cilium-localizing proteins in three ciliated cells of diverse origin. We reveal that the primary cilium shows the most heterogeneous proteome compared to all other organelles, across cell types and within the same cell population, and that the heterogeneity relates to cell-individual customization of the cilium to conduct cell-specific sensory signaling. Furthermore, we demonstrate that our cilia atlas paves new avenues for cilia and ciliopathy research, by leveraging its content for relating ciliary protein localization to the cell cycle, revealing a new cilia marker and identifying a potential novel ciliopathy gene.

## Results

### Mapping the subcellular and subciliary location for human proteins

To examine the proteomic composition of primary cilia on the single-cilium level, we developed an antibody-based spatial proteomics workflow, which applies the antibody library of the Human Protein Atlas (HPA) ^27,28^ together with markers for nuclei, cilia, and basal bodies (Figure 1A) and high-throughput, high-resolution 3D confocal microscopy (Figure 1B). With this workflow we comparatively analyzed three immortalized primary human cell lines of different tissue origin and sex: (1) serum-starved hTERT-RPE1 cells from the embryonic retina ^29^, to represent the best studied *in vitro* model for human cilia, (2) RPTEC/TERT1 cells ^30^, derived from the epithelial kidney proximal tubules, to represent a cell type affected in the most common ciliopathy - polycystic kidney disease (PKD), and (3) the mesenchymal, multipotent, stromal ASC52telo cells ^31^, derived from adipose tissue, to represent a multipotent cell type. The three cell lines are morphologically diverse and likely rely on different signaling and ciliogenesis programs, allowing coverage of a broad band of cellular functionality and ciliary diversity (Figure 1C). The starved hTERT-RPE1 cells represent a synchronized culture with a high ciliation rate and that generates cilia mostly at the cell-glass-interface (Figure 1D). Reportedly, hTERT-RPE1 cells predominantly employ the internal ciliogenesis pathway ^5^. The RPTEC/TERT1 cell culture generates polarized cells that cycle slowly and grow long cilia sticking into the fluid (Figure 1D). In contrast, the ASC52telo cells cycle fast and present short, flat-lying cilia (Figure 1D). We observed ASC52telo cells to occasionally show dots or small lines of the ciliary marker on one centrosome during mitosis, hinting to the retainment of a ciliary vesicle or cilium during mitosis, similar to reported observations in HEK293T cells ^32^ (Figure 1E). This was not observed in the other cell lines. However, in RPTEC/TERT1 cells, we occasionally observed cells that formed two adjacent cilia (Figure 1F). Double ciliation was also seen in hTERT-RPE1 cells, albeit much rarer (Figure 1G). Noteworthy, we once observed a potential ciliary bridge in RPTEC/TERT1 cells, where cilia from neighboring cells appeared to be either connected or touching (Figure 1H), as previously described for murine cells ^33^.

**Figure 1:**
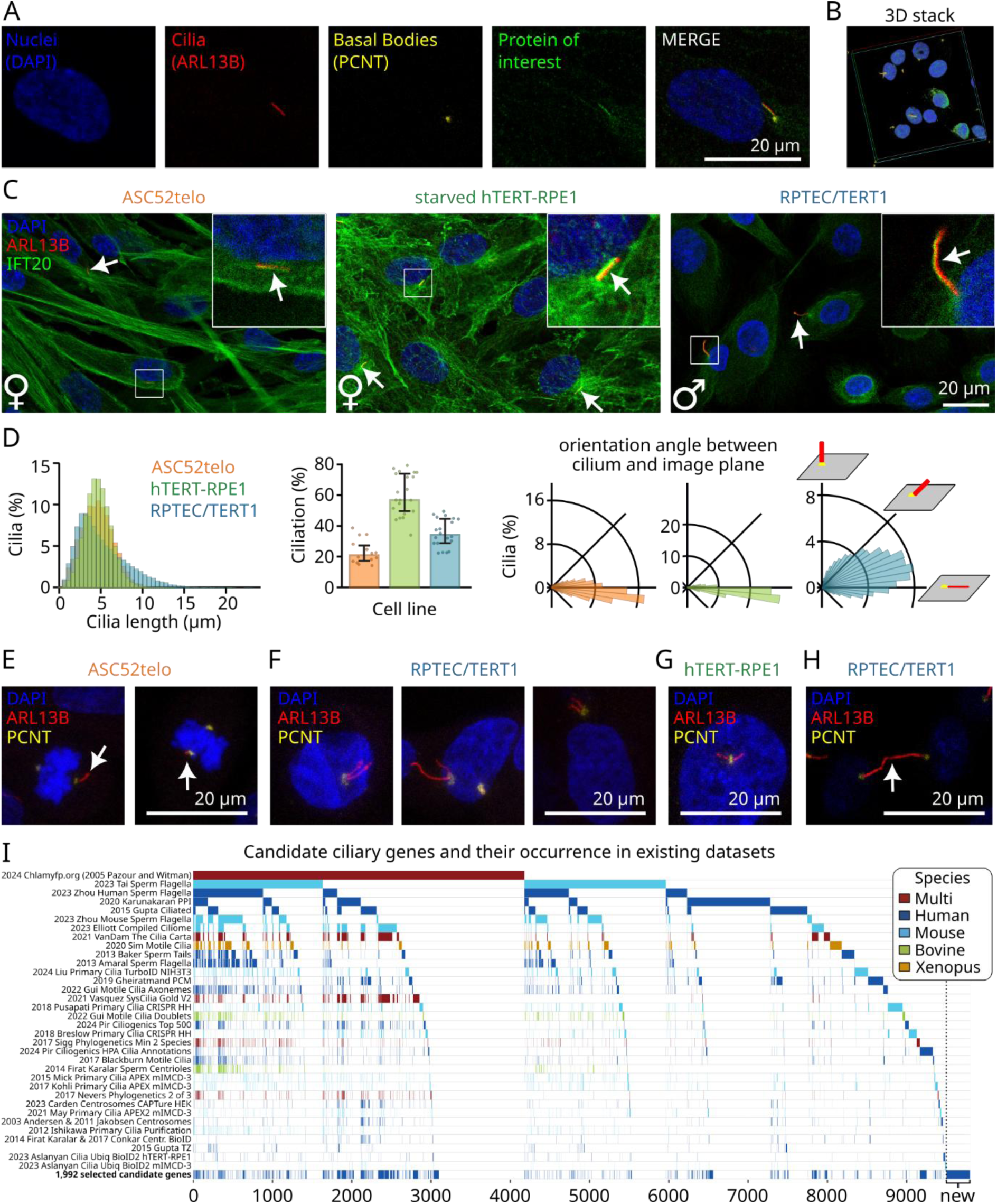
Spatially profiling the primary cilia proteome. **A-B)** New staining concept. **A)** Ciliated cells (i.e., starved hTERT-RPE1) cells are co-stained with markers for nuclei (DAPI), cilia (ARL13B antibody), basal bodies (PCNT antibody) and for a protein of interest (i.e., with a polyclonal antibody against IFT43 from the Human Protein Atlas). **B)** Confocal z-stacks are generated, which allow 3D analysis of the whole cell including the cilia, even when they stick out of the cell to the top or bottom (Blue: Nuclei, Red: Cilia, Yellow: Basal Body, Green: Protein of Interest). A 3D projection of an example z-stack is shown. **C)** Three cell lines are investigated: ASC52telo cells from the adipose tissue, starved hTERT-RPE1 cells from the retina, and RPTEC/TERT1 cells from the kidney. Example images from all three cell lines are shown with magnified insets, from a co-staining of DAPI (labeling nuclei), ARL13B (labeling cilia), and the protein of interest IFT20. Gender of the cell line indicated. **D)** CiliaQ analysis of all images underlying this study reveals the cilia length distribution, ciliation rate, and ciliary orientation angle. Percentages of cilia refer to the total number of cilia by cell line. In total 280,515 cells with 128,156 cilia were included. Left plot: Normalized histograms of the cilia lengths for all cilia analyzed, distinguished by cell line. Middle plot: Each dot represents all images from one individual experiment day. Bars and error bars represent median and quantiles, respectively. Right plot: Radial histograms of the ciliary orientation angles for all cilia analyzed. The orientation angle was determined as the angle between the vectors from the basal body center to the ciliary tip and the image plane. **E-H)** Example images of special cilia observations in the three different cell lines. Only the nucleus (DAPI), cilia (ARL13B), and basal body (PCNT) channels are displayed. **E)** Mitotic ASC52telo cells retaining a cilium. **F)** RPTEC/TERT1 cells with two adjacent cilia. **G)** Starved hTERT-RPE1 cell with two adjacent cilia. **H)** Two adjacent RPTEC/TERT1 cells whose cilia are connected and form a “bridge”. **I)** Matrix plot of the different available cilia data sets accumulated to form the cilia candidate list of 9808 human genes demonstrating the heterogeneity of existing cilia datasets and their coverage of the candidate proteins (see STAR Methods). Each study is represented by one row, the presence of a gene in the study is marked by color. The color indicates the species the study was conducted in. The bottom row marks the 1,992 candidate genes selected for staining with Human Protein Atlas antibodies in this study, including 298 educated guesses of candidate genes that had not been associated with cilia before (“new”). Data available in Table S1.

For the targeted antibody-based spatial proteomics approach, we assembled a list of candidate proteins based on existing curated databases including SYSCILIA gold standard version 2 ^34^ and the ChlamyFP.org database ^14^, the tissue resource of the HPA ^27,35^, MS data sets on primary cilia based on proximity-labeling approaches or purification attempts ^20–25,36^, predictive studies including the CiliaCarta ^37–39^, comparative transcriptomics studies ^40^, phylogenetic studies on flagellated species ^41,42^, CRISPR screens ^43,44^, MS studies of flagella or motile cilia ^45–49^, Cryo-ET studies of sperm flagella ^50–52^, and studies of the centrosomal proteome ^53–58^. For a strong focus on signaling pathways, we extended the screen with key components of broad signaling pathways, such as AKT kinases, MAP kinases, mTOR, a selection of G-protein coupled receptors (GPCRs), phosphodiesterases (PDEs), or adenylyl cyclases, even without any prior evidence of localizing to cilia. Interestingly, our assembled cilia candidate list highlighted that nearly ten thousand protein-coding genes have been reported or suggested to potentially produce proteins localizing to primary cilia or engaging in their regulation (Figure 1I, Table S1). The overlap of proteins identified in cilia with different methods or in different cell types or species was particularly small: 3760, 1736, and 1017 genes were attributed to cilia in one, two, or three data sets only, respectively (Figure 1I). This may point to an incredible heterogeneity in the ciliary proteome across species and cell types, and/or potentially many false positives due to technical limitations. Finally, we prioritized the candidate list and selected 1,992 candidate proteins for assessment of ciliary localization in human cells (see Materials and Methods) (Figure 1I, Table S1).

### A spatial map of the human cilia proteome

Applying our spatial proteomics data generation pipeline targeting 1,992 proteins in ciliated cells produced a massive high-resolution 3D confocal image data set of cilia and their protein composition with 19,905 3D image stacks, revealing 280,515 cells, 128,156 of which displayed a cilium. We applied the established HPA subcellular workflow to annotate all images, which involves investigation and review of every staining by multiple skilled cell biologists ^28^, and provide example 3D images and annotations for all stainings passing quality control on the webpage of the HPA (https://proteinatlas.org) where they can be interactively explored. In addition to the 35 subcellular organelles and structures annotated in the HPA, we annotated protein localization to the primary cilium, basal body, transition zone, and ciliary tip (Figure 2A and Figure 2B). We could identify the proteins encoded by 715 of our 1,992 candidate genes at cilia, of which 378 proteins were detected in primary cilia, 117 at the ciliary tip, 100 at the transition zone region, and 438 at the basal bodies (Figure 2C). Furthermore, we observed additional subpatterns of ciliary localization, such as partial staining of the proximal cilium (e.g., for EFCAB7, EVI2A, and TRIM46, Figure 2D), which might correspond to a domain distal from the transition zone and close to the “inversin” compartment ^59^, such as the EVC-EVC2 complex described in mouse fibroblasts ^60^, or a staining of the ciliary pocket; spotty staining patterns along the cilium (e.g., for TRIO, Figure 2D); or enrichment towards the ciliary tip (e.g., for C2CD4C, Figure 2D); or combinations of spotty patterns and enrichment towards the tip for the same protein (e.g., for TMEM237, Figure 2D), highlighting the subciliary resolution of our screen.

**Figure 2:**
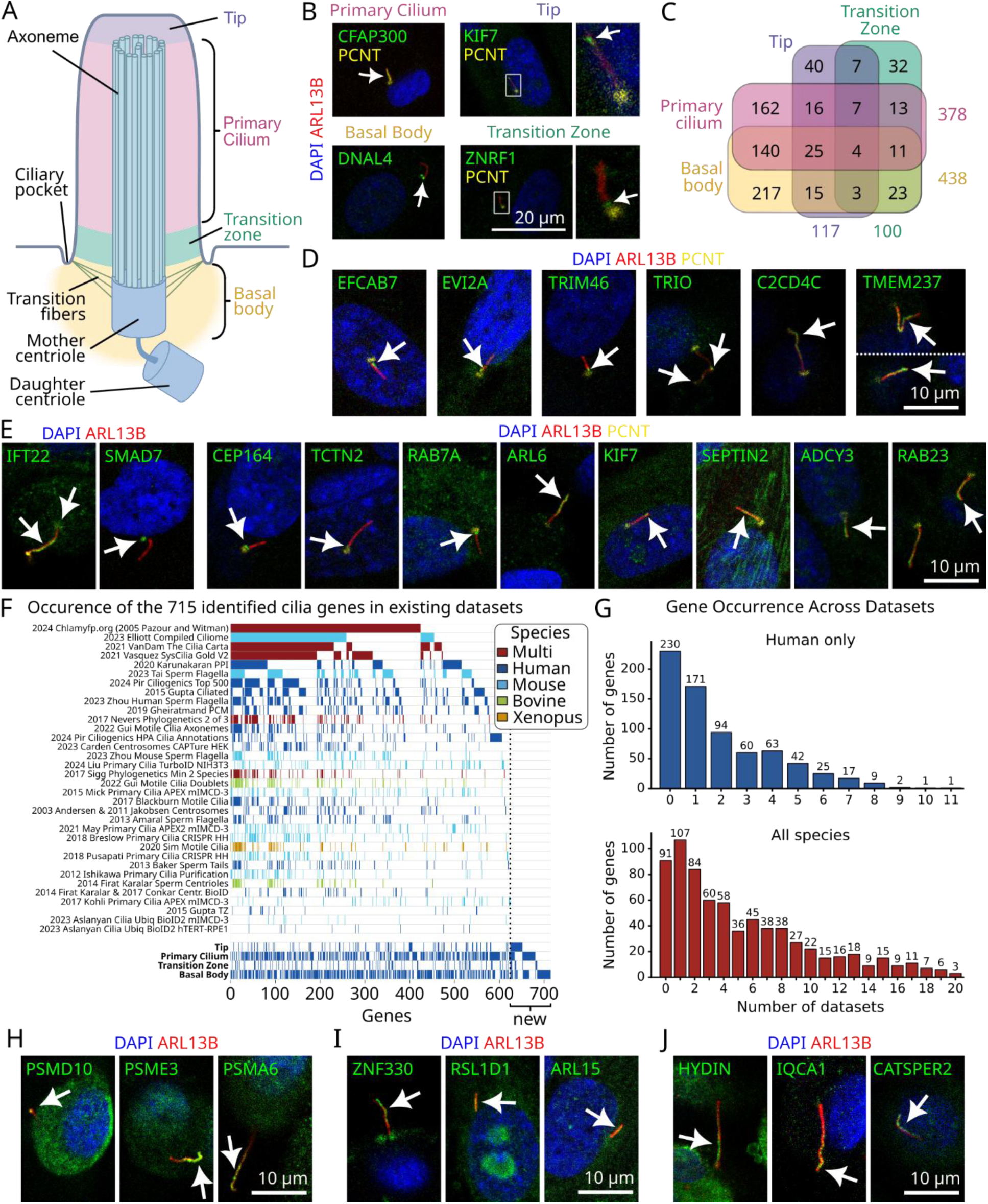
Spatial map of the primary cilia proteome. **A)** Schematic drawing of a primary cilium and its substructures. **B)** Protein localization to four ciliary substructures was annotated, for which example images are shown: Primary cilium, if the protein was overlapping with the ciliary marker ARL13B; Primary cilium tip (Tip), if the protein of interest channel showed a signal at the distal end of the cilium; Basal body, if the protein of interest channel showed signals overlapping or accumulating close to the basal body marker PCNT; Primary cilium transition zone (Transition zone) if the protein of interest channel showed an enrichment between basal body and cilium. **C)** Venn diagram showing the number of genes observed at one or multiple of the four locations specified in B. **D)** Example images of additional subpatterns observed in primary cilia. Gray interrupted line distinguishes two individual images placed adjacently for showing two cilia. **E)** Example images of stainings for proteins that are established as localizing to cilia - gene names are shown. **F-G)** Comparing the list of genes encoding proteins that were annotated as localizing to at least one of the four cilium locations (B) to existing datasets. **F**) Matrix plot showing each identified gene in a single column, whether it occurred in a data set (denoted by color, color encodes data set species, data sets in upper rows), and where it was identified in this study (bottom rows, bold labels). 91 “new” proteins were not found in any other dataset. Data available in Table S1. **G)** Replotting the matrix (F) as a histogram showing how many genes (y axis) occurred how many times (x axis) in other data sets, considering only data sets restricted to human cilia (top) or cilia on any species (bottom). **H-J)** Example images of proteasomal subunits observed in cilia (H), proteins observed in cilia that were previously only detected in individual proximity-labeling-based MS approaches (I), and motile cilia proteins detected in the cilia of RPTEC/TERT1 cells. Images were derived from stainings of different cell lines: Starved hTERT-RPE1 cells: DNAL4, ZNRF1, EFCAB7, EVI2A, TRIM46, SMAD7, RAB7A, SEPTIN2, RAB23, ARL15; RPTEC/TERT1 cells: CFAP300, TRIO, C2CD4C, TMEM237, IFT22, ARL6, PSMD10, PSME3, PSMA6, HYDIN, IQCA1, CATSPER2; ASC52telo cells: KIF7, CEP164, TCTN2, KIF7, ADCY3, ZNF330, RSL1D1. Single confocal slices extracted from a 3D stack are shown in the images.

Among the revealed proteins are many established ciliary proteins, such as the intraflagellar transport protein IFT22, SMAD7 localizing to the basal body, the subdistal appendage protein CEP164, the MKS-complex protein TCTN2 at the transition zone, the small GTPase RAB7A in the transition zone, the ciliary trafficking protein ARL6 enriched towards the ciliary tip, the kinesin KIF7 at the ciliary tip, the septin SEPTIN2, the adenylyl cyclase ADCY3, or the small GTPases RAB23, which is known to participate in regulating GLI transcription factors and ciliary Hedgehog signaling (Figure 2E). ^61,62^ This confirms the validity of our approach for diverse types of ciliary proteins and provides the cilia field with subcellular resolution pictures of these and many more known proteins in unperturbed human cells. In total, we identified 91 proteins that had never been previously associated with any type of cilium across different species (Figure 2F, Table S1). Furthermore, 230 proteins had never been identified to localize to human cilia by MS- or Cryo-ET based approaches (Figure 2G). Notably, our image-based screen complimentarily extends our knowledge on proteins previously only detected with MS in cilia - we validate at least 37 proteins previously listed in MS datasets only (Table S1) and provide their subciliary localization patterns.

An interesting new discovery were three proteasomal subunits in cilia: PSMD10, PSME3, and PSMA6 (Figure 2H). PSMD10 is among the 91 proteins never related to cilia before. PSME3 has previously been detected in one mouse sperm flagella data set ^50^ and was bioinformatically predicted to be a cilia protein ^37^. PSMA6 was identified in cilia by many approaches and was listed in databases ^14^, an analysis of murine kidney cilia with cilia-APEX ^22^, a CRISPR-based screen ^44^, a comparative transcriptomic analysis ^40^, and in motile cilia and sperm flagella studies ^45,48,50–52^. Other examples for our data confirming proximity-labeling-MS-approaches in the cilia of model organisms are ZNF330 or the senescence-regulator RSL1D1 (Figure 2I), which both had previously been identified by cilia-APEX in murine kidney cells ^21,22^. Also, we confirm previous bioinformatic predictions, such as ARL15 in cilia (Figure 2I) ^39^. Some of the studies in our candidate list analysis were from motile cilia. Surprisingly, we identified proteins believed to be specific for motile cilia in primary cilia of RPTEC/TERT1 cells, such as the central-pair protein HYDIN, IQCA1, which is part of the nexin-dynein-regulatory complex in motile cilia, or the CATSPER2 channel subunit (Figure 2J). This is particularly interesting in light of recent descriptions of motile cilia proteins and active motility of primary cilia in *ex vivo* pancreatic islet cells ^63^. Yet, we see that HYDIN, IQCA1, and CATSPER2 accumulate towards the tip rather than along the whole cilium, which might indicate for HYDIN and IQCA1 that they are not in fact implemented into the axoneme but might just transit the cilium. Of note CATSPER2 is expressed in RPTEC/TERT1 cells according to the HPA cell line sequencing data confirming staining validity. Further studies will need to delineate why and to which purpose motile cilia proteins transit RPTEC/TERT1 cilia and whether this finding translates to primary cilia of proximal tubule cells *in vivo*. However, kidney proximal tubule cells of patients with kidney injury have been observed to feature multi-ciliated cells whose cilia ultra-structurally show features of motile cilia ^64^, suggesting that proximal tubule cells may in fact be able to establish ciliary motility.

### The primary cilium is a hub for diverse sensory signaling pathways

To explore the functions of the identified cilia proteins, we performed an enrichment analysis for Gene Ontology (GO) Biological Process terms (Figure S1A and Figure S1B, Table S2). Beyond many terms relating to maintenance processes of the cilium or basal body (e.g. intraciliary transport, adj. p≤1.5e-27 and cilium organization, adj. p≤1.3e-93), we found significant enrichment of terms relating to signaling, such as calcium signaling (calcium-mediated signaling, adj. p=9.5e-06 for primary cilium), Wnt signaling (canonical Wnt signaling pathway, adj. p≤3.1e-05), and morphogenesis and developmental signaling pathways (e.g. regulation of developmental growth adj. p≤0.0034) (Figure 3A, Table S2). In fact, we detected signaling proteins of most major signaling pathways in the primary cilium (Table 1) of which many had hitherto not been confirmed in cilia. For example, we revealed in cilia the kinases AKT1 and AKT3, the Serine/threonine-protein kinase TAO1 (TAOK1), a catalytic and regulatory subunit of PI3K (PIK3CA and PIK3R4), the focal adhesion kinase PTK2, the calcium/calmodulin dependent protein kinase CAMK1G, the protein phosphatase 2 (PPP2R3C), the E3 ubiquitin-protein ligase MIB1, the vascular endothelial growth factor receptor 3 FLT4, the GPCR OPRL1, the src-related receptor PDGFRA, the transforming growth factor beta receptor 1 TGFBR1, and the cyclic nucleotide gated ion channel CNGA2 (Figure 3B). Of note, beyond the GPCR OPRL1, we revealed several additional GPCRs in cilia using HPA antibodies that have been recently confirmed to be highly selective for their respective target ^65^, such as OPN3, CHRM3, or CXCR2, extending the list of described cilia-localizing GPCRs ^1^ further (Table 1). Another example for a ciliary signaling protein identified is the Triple functional domain protein TRIO mentioned above (Figure 2D), a Guanine nucleotide exchange factor (GEF) for RHOA and RAC1 GTPases ^66^ and which reportedly mediates signal transduction in the recently discovered axon-cilium-synapse ^67^. In summary, these results highlight the cilium as a versatile signaling organelle with broad signaling capacities, filled with most receptor types and diverse downstream signaling machinery.

**Figure 3:**
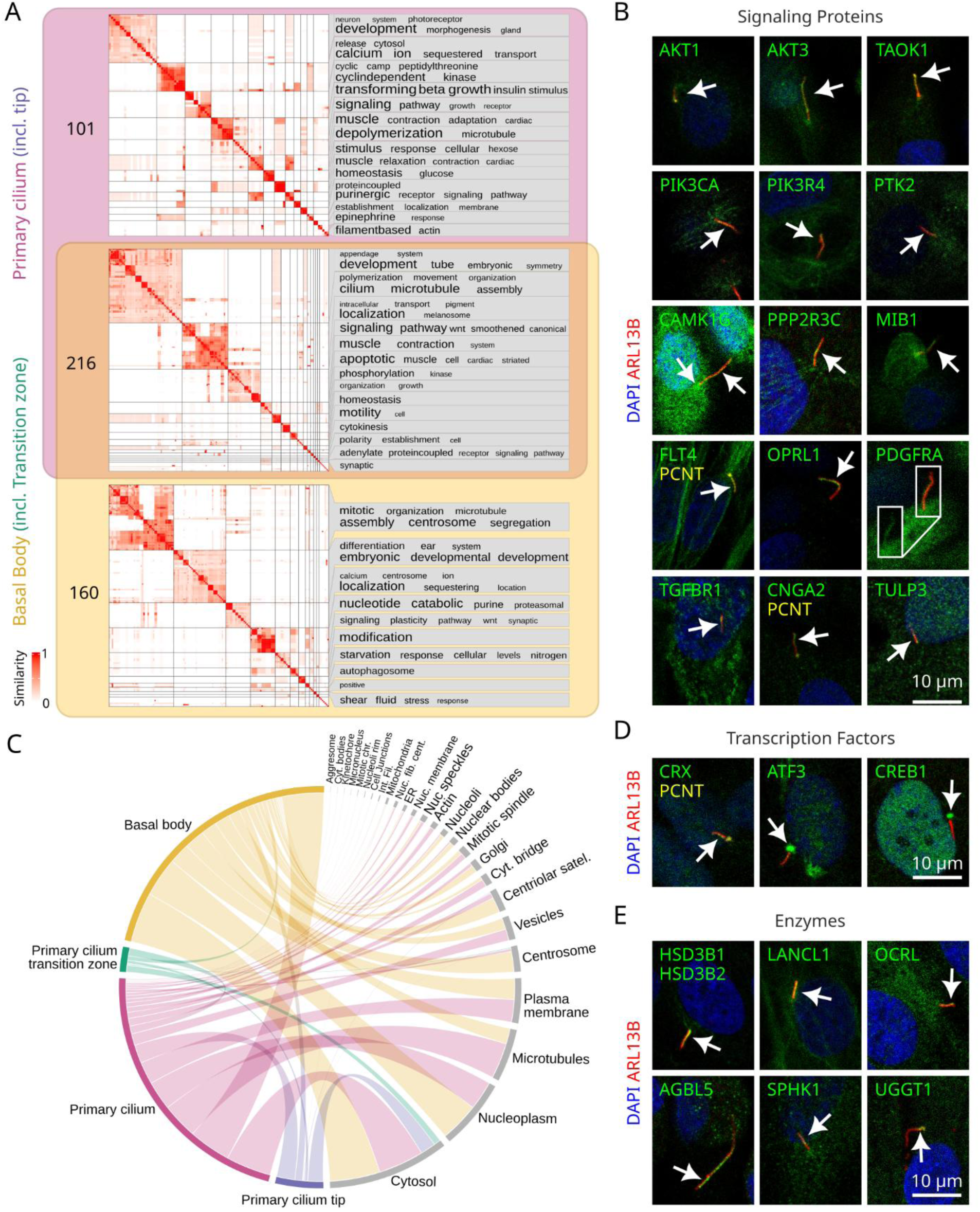
The primary cilium is a signaling hub. **A)** Comparative Gene Ontology enrichment analysis of Biological Process terms between the two major cilia locations primary cilium (including tip) and basal body (including transition zone). The Venn diagram shows the number of terms found to be significantly enriched (adj. P value < 0.01 and q value < 0.01) for only primary cilium (n=125) or basal body (n=129), as well as for both locations (n=198). For significant terms the pairwise semantic similarities were calculated and subsequently clustered (heatmap). For each resulting GO term cluster, a word cloud is shown summarizing the biological functions of the terms in the cluster based on keywords enrichment of term names. The font size in the word clouds correlates with -log(P value) of the key word enrichment, i.e. the larger, the more enriched a keyword is compared to background GO vocabulary. The color in the heat maps represents the semantic similarity. **B)** Protein localization of example signaling proteins to the primary cilium. All images show single confocal slice images except for PDGFRA, where a maximum intensity projection of five confocal slices is used to show the whole cilium. The inset for PDGFRA shows the same position with displaying only the PDGFRA channel. **C)** Chord diagram of protein multi-localization to the different subcellular compartments for all proteins localizing to at least one of the ciliary compartments. Each observed localization to a ciliary compartment and another denoted compartment in the same image is contributing to the thickness of the lines. Abbreviations: Centriolar Sat.es = Centriolar Satellites; Cyt. bridge = cytokinetic bridge; Int. Fil. = Intermediate Filaments; Mitotic chr. = Mitotic chromosome; Nuc. bodies = Nuclear bodies; Nucleoli fib. cent. = Nucleoli fibrillar center; Nuc. Membrane = Nuclear membrane; Nuc. speckles = Nuclear speckles. **D)** Example transcription factors revealed in cilium and nucleus. **E)** Example enzymes revealed in cilia. Images were derived from stainings of different cell lines: Starved hTERT-RPE1 cells: PTK2, PP2R3C, CNGA2, TULP3, CRX, ATF3, CREB1, HSD3B1/HSD3B2, LANCL1, OCRL, SPK1; RPTEC/TERT1 cells: AKT1, AKT3, TAOK1, PIK3CA, PIK3R4, CAMK1G, MIB1, OPRL1, PDGFRA, AGBL5, UGGT1; ASC52telo cells: FLT4, TGFBR1.

**Table 1.**
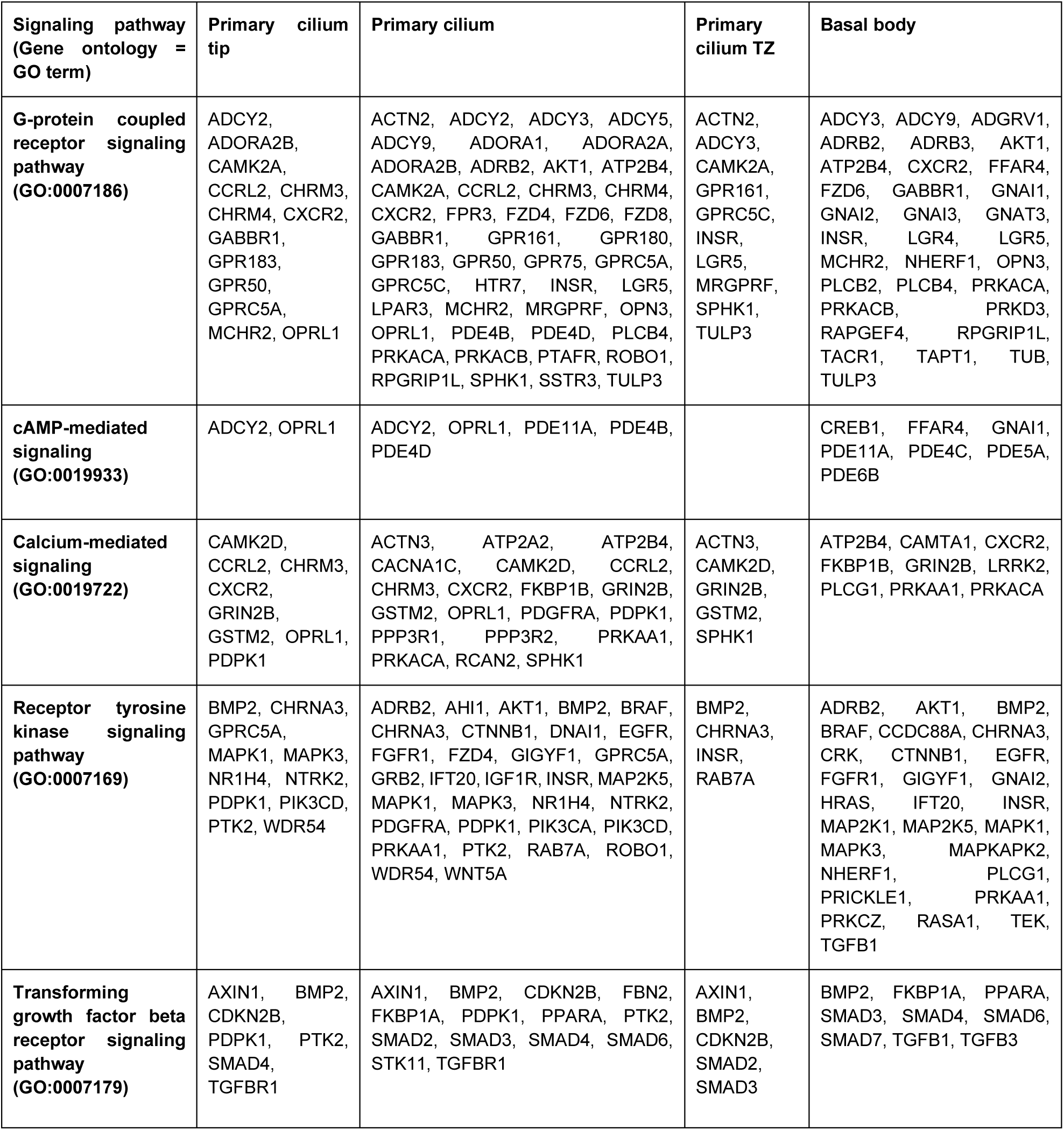
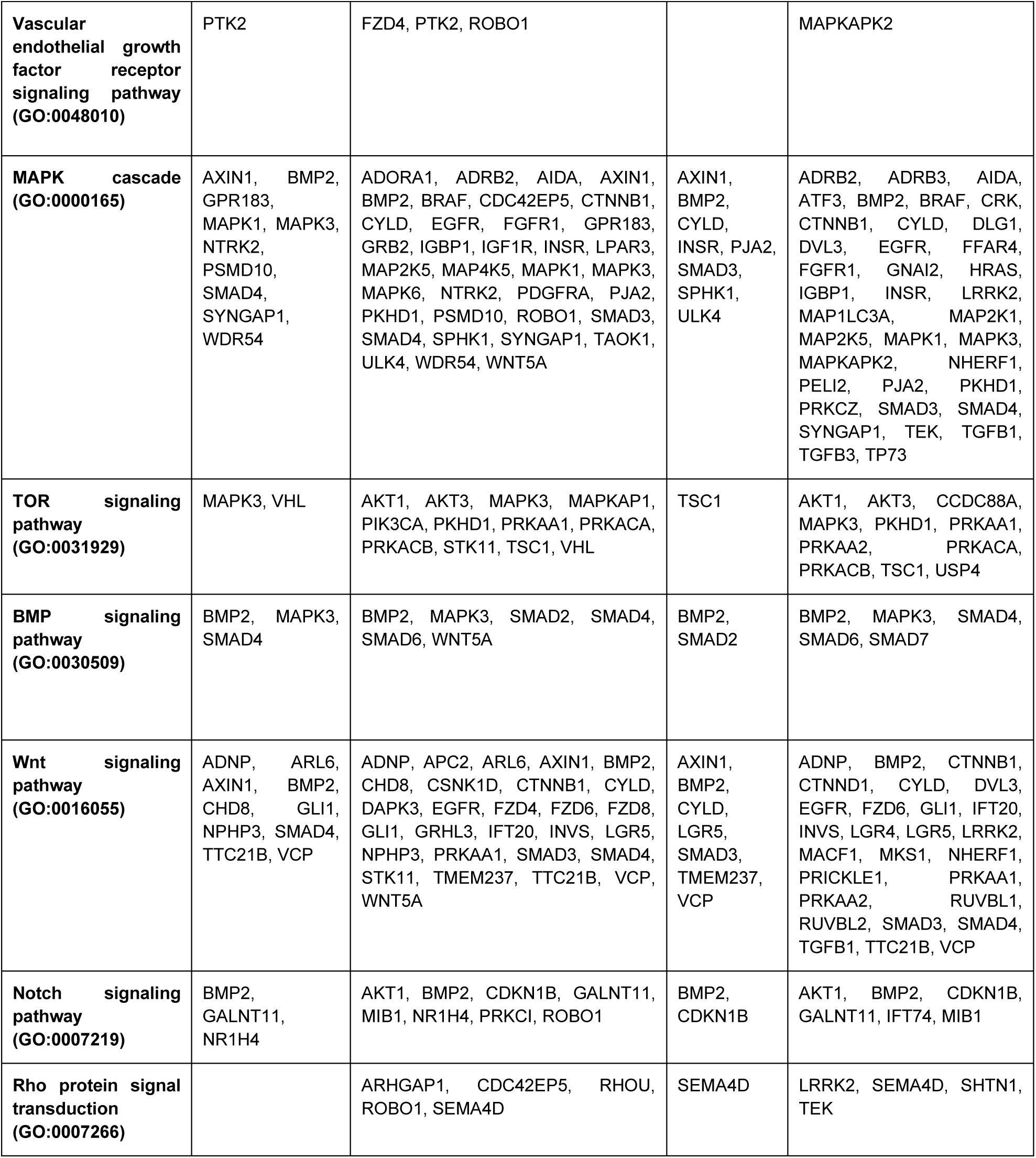
Signaling proteins revealed in cilia or basal bodies. Gene names part of the selected GO terms are shown if they were revealed at the primary cilium tip, primary cilium, primary cilium transition zone (TZ), or the basal body.

### Subcellular protein multilocalization and the implications for ciliary function

Protein localization is in general closely associated with protein function. We and others have previously shown that more than half of the human proteome is localized to multiple subcellular compartments, i.e. are multilocalizing ^28,68,69^. Here we show that cilium-localizing proteins (including tip and transition zone) are significantly more multilocalizing than other proteins (p=3.8e-202) - as much as 99% of cilium-localizing proteins are also found in other cellular compartments (Figure S1C). Looking at individual stainings, we observed that the protein was multilocalizing to a ciliary and another subcellular compartment in the same cell culture population in 92.5% of the cases, with the most common other compartments being basal bodies, cytosol, nucleoplasm, and microtubules (Figure 3C). In light of the basal body being the access point to the cilium ^10^, it makes sense that we observed many ciliary proteins also at the basal bodies, potentially transiting the basal body region during transport to cilia. Multilocalization with microtubules may, to some extent, relate to proteins that interact with the microtubules in the ciliary axoneme as well as with cytoplasmic microtubules, such as tubulin modifying enzymes like TTLL6 or TTLL9.

In contrast, proteins localizing at cilia and in the cytosol and nucleoplasm may rather be proteins involved in relaying information from the cilium to the cell body or potentially engaging in similar or different functions or signaling pathways in different places in the cell (a process called “moonlighting” ^70^). In fact, we identified kinases that are central in many signaling pathways in cilia, but also in cell body compartments, i.e., the cytosol, such as AKT1 and AKT3, PIK3CA and PIK3R4, or PTK2 (Figure 3B). Their ability to localize to multiple locations may be attributed to their capacity to transduce distinct signals in various cellular compartments, influenced by the diverse environments with varying stimulators and substrates. We also observed proteins with different ciliary and cell body functions in multiple locations in the cell. For example, the protein TULP3 is well known for a role as a cargo-adaptor in IFT-A mediated trafficking of GPCRs and ARL13B to the cilium ^71–73^, but may also act as a transcription factor residing at the plasma membrane and relocalizing to the nucleus upon activation ^74^. In agreement, we detected TULP3 in cilia, at the plasma membrane, and in the nucleus (Figure 3B). Many observations of multilocalization for cilia proteins may also relate to the cilium relaying information to the cell body. For example, for Gli transcription factors in Hedgehog signaling it is established that post-translational modification of the transcription factors occurs in the cilium, upon which they are transported into the cytosol, where they are - depending on the modification - either degraded or sent to the nucleus to adjust gene transcription ^1^. We in fact found additional transcription factors at cilia, including CRX in cilia (Figure 3D), or ATF3 and CREB1 at the basal body (Figure 3D), which co-localized to the nucleoplasm, supporting the hypothesis that the cilium controls gene expression by local modification of transcription factors beyond Sonic Hedgehog signaling, as shown for CREB1 ^75^.

Another class of proteins that often multilocalize and “moonlight” are enzymes, as we recently demonstrated ^76^. Little is known about metabolic and enzymatic activities in the cilium. Notably, not only the proteomic but also the lipid composition of the cilium makes up the unique ciliary compartment ^77^. While the lipid composition of cilia has been shown to be unique compared to the plasma membrane, the processes involved in its formation - whether through transport mechanisms or local production - remain underexplored. While for phosphoinositides it has been shown that the membrane composition is locally regulated by enzymes, for other lipid species it is not understood how they end up in the ciliary membrane, such as for cholesterol, which is described to essentially regulate Hedgehog signaling ^77,78^. Interestingly, we discovered several enzymes in cilia, which also localize to other positions in the cell. For example, we detected an enzyme that is part of sterol synthesis in cilia (HSD3B1/HSD3B2, Figure 3E). Other examples are the Glutathione S-transferase LANCL1, the Inositol polyphosphate 5-phosphatase OCRL, the carboxypeptidase AGBL5 that is involved in tubulin de-glutamylation and has been indicated as a negative regulator of ciliogenesis ^79^, the Sphingosine kinase SPHK1, which regulates ceramide levels and is highly engaged in signaling ^80–82^, or the UDP-glucose glycoprotein glucosyltransferase 1 (UGGT1), a protein that reportedly acts as a gate-keeper for folding of glycosylated proteins at the endoplasmic reticulum (ER) (Figure 3E) ^83,84^.

Our data demonstrates that cilia mainly contain multilocalizing proteins and reveal several new enzymes in cilia, supporting the hypothesis that the ciliary lipid composition is shaped and adjusted locally through enzymatic activity and highlighting the capabilities of the cilium to modify molecules and proteins locally.

### Cilia proteomes reveal cell type specific functional differences

Most of the investigated proteins (69%) localized to cilia only in one or two of the studied cell lines (Figure 4A). In the cell lines where we did not observe ciliary localization, the protein was not detectable or localized to other compartments. For example, we identified the calmodulin- and kinase-related signaling protein STRN3 at the plasma membrane / actin cytoskeleton in all three cell lines, but only in RPTEC/TERT1 cells in the cilium and we identified the common kinase PI3K (PIK3CA) in cilia in RPTEC/TERT1 and hTERT-RPE1 cells but not in ASC52telo cells (Figure 4B). Other examples are the activin receptor ACVR1B, the SEPTIN9, the RHO GTPase activating protein ARHGAP1, or the adenylyl cyclase ADCY3. We observed ADCY3 to be more enriched in cilia of ASC52telo cells than of RPTEC/TERT1 cells, while it co-localized to the plasma membrane in both cell lines. In hTERT-RPE-1 cells, ADCY3 was confined to basal bodies, similar to a previous report ^85^ (Figure 4B). Such cell-type specificity of ciliary protein localization is higher than for other organelles, suggesting that the ciliary proteome is more tunable than the proteome of other organelles and compartments.

**Figure 4:**
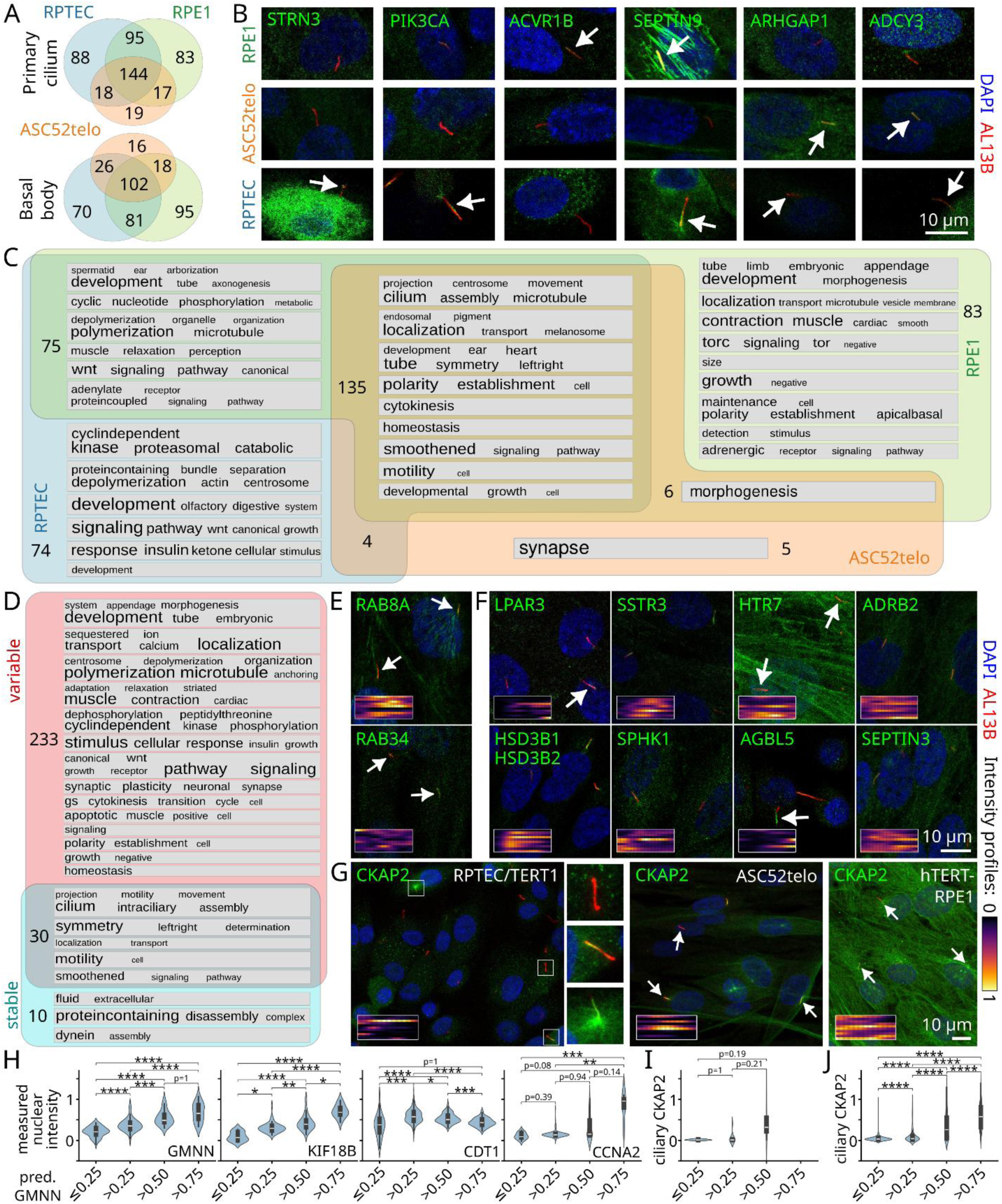
Cellular diversity and spatiotemporal dynamics of the primary cilia proteome. **A)** Venn diagram of the genes for the proteins identified in primary cilia (including tip and transition zone), distinguishing by cell line, only including genes, for which an approved staining was obtained from all three cell lines. (Abbreviations: RPTEC = RPTEC/TERT1, RPE1 = starved hTERT-RPE1) **B)** Example images (single confocal slice images) for stainings across the three cell lines, showing heterogeneity in ciliary localization across cell types. **C)** Comparative Gene Ontology enrichment analysis of Biological Process terms between the three included cell lines to investigate cell-line differences and commonalities. The Venn diagram shows the overlap of significantly enriched terms (adj. P value < 0.01 and q value < 0.01) for each cell line. Significant terms for each section of the Venn diagram were clustered by semantic similarity, and for each resulting GO term cluster, a word cloud is shown summarizing the biological functions of the terms in the cluster based on keywords enrichment of term names. The font size in the word clouds correlates with -log(P value) of the key word enrichment, i.e. the larger, the more enriched a keyword is compared against background GO vocabulary. Heat maps displaying the semantic similarities are shown in Figure S3. **D)** Comparative Gene Ontology enrichment analysis of Biological Process terms between proteins annotated as displaying intensity variation in one of the four ciliary locations, and annotated as stable as described in C). Heat maps displaying the semantic similarities are shown in Figure S5. **E)** Example images (single confocal slice images) for stainings of the two ciliogenesis-related proteins RAB8A and RAB34. **F)** Example images (single confocal slice images) of stainings revealing proteins with heterogeneous localization to primary cilia. **G)** Example images (maximum intensity projections of whole stack) of CKAP2 staining in the three cell lines. Cell lines for images (if not indicated): Starved hTERT-RPE1 cells: LPAR3, SSTR3, HTR7, ADRB2, HSD3B1 HSD3B2, SPHK1, RAB8A, SEPTIN3; RPTEC/TERT1 cells: AGBL5; ASC52telo cells RAB34. Insets in E-G show the intensity profiles of the proteins of interest for 10 individual example cilia (each row represents one cilium) from the center of the basal body (left) to the tip (right), revealed by CiliaQ analysis (see Materials and Methods). **H)** Validation of a GMNN prediction model for 3D image stacks of RPTEC/TERT1 cells, applicable for estimating the cell cycle state (x axis: predicted nuclear average GMNN intensity, y axis: measured nuclear intensity of the different proteins indicated, revealed by antibody staining). **I,J)** Relationship of ciliary CKAP2 intensity (average intensity of centerline) and predicted cell cycle state, determined from the four CKAP2 image stacks recorded in the cilia atlas yielding 91 cells (I) or from a data set with 150 3D stacks acquired for validation yielding 4,149 cells (J). Statistical tests: Krusal-Wallis with Dunn’s test for multiple comparison correction. Significant p-values are indicated as * for p ≤ 0.05, ** for p ≤ 0.01, *** for p ≤ 0.001, **** for p ≤ 0.0001.

In a comparative GO term enrichment analysis of the identified cilia proteins between cell lines (Figures 4C and S2A, Table S3) we observed that indeed each cell line has some unique ciliary specializations. While all cell lines shared GO terms relating to general cilia processes such as (1) ciliogenesis, microtubules, and anchoring of the cilium, (2) transport processes, (3) smoothened signaling, (4) development, and (5) cell polarity (Figures 4C, S2B, S2C, S2D, and S2E), each cell line was also associated to unique GO terms: RPTEC/TERT1 cilia appeared to be tailored to various sensory signaling pathways whereas hTERT-RPE1 cilia appeared to be more focused on development and morphogenesis, and ASC52telo cilia displayed proteins related to synapse organization and glutathione metabolism. Since this study is limited to three cell lines, we expect this heterogeneity to be even higher when extending to more cell types.

### Single cilia heterogeneity and its relation to cell states

Thanks to the single cell resolution of our imaging-based approach, we were able to assess variations in protein localization and abundance within primary cilia at the individual cell level. Beyond heterogeneity across cilia in different cell types, we also observed an incredible heterogeneity on the single cell level. 389 of 498 (78%) of the primary cilium, primary cilium tip, or primary cilium transition zone stainings were classified as varying across individual cells. Using a comparative GO term enrichment analysis, we explored which processes are particularly stable and which are particularly variable across individual cilia. As expected, proteins engaging in ciliogenesis, microtubules, and ciliary transport-associated processes appeared in both groups, stable and variable (Figures 4D and S3, Table S4), while the most dynamic localization was observed for proteins involved in signaling, developmental, and morphogenesis processes. As an example for ciliogenesis-related single-cell heterogeneity, RAB8A and RAB34, which are both known to shape the cilium during ciliogenesis by mediating transport of proteins to the growing cilium, localized only to individual cilia (Figure 4E). Interestingly, some GPCRs showed varying abundance levels across cilia even in the serum-starved, synchronized hTERT-RPE1 cells - e.g., LPAR3, SSTR3, and HTR7 largely varied in ciliary abundance whereas ADRB2 appeared more stable (Figure 4F). Furthermore, many enzymes in cilia showed single-cilia variation, such as for HSD3B1/HSD3B2, SPHK1, and AGBL5 (Figure 4F). Besides locally contributing to shaping a specific lipid environment, such enzymes may also engage in the process of lipid-dependent regulation of signaling, such as for Hedgehog signaling, which has been shown to be regulated by cholesterol ^78,86^. Single-cilia variation was also seen for Septins, such as for SEPTIN3, which we newly identified in cilia and which heterogeneously localized to hTERT-RPE1 cilia but not to cilia of the other cell lines (Figure 4F). Septins are known to be cell type specific and attach to the membrane in a PIP-dependent manner ^87^. Observing Septin heterogeneity in single cilia hints at a heterogeneous nature of ciliary lipid-protein-interactions. In summary, variation in ciliary localization of GPCRs and lipid-metabolic or -interacting enzymes may relate to the signaling activity of cilia or to fine-tuning of the ciliary signaling machinery.

For some proteins, we observed the variation to correlate with localization of the protein to other cell compartments, such as for CKAP2 in RPTEC/TERT1 cells, where localization to the cilium and part of the microtubule cytoskeleton were seen in some cells, and localization to vesicles in other cells (Figure 4G). Given that CKAP2 was known to be related to the cell cycle, we inferred that ciliary localization might be related to the cell cycle as well. Visually, postmitotic cells showed high intensity for the vesicle staining. A possible transitional stage where the cilium showed staining for CKAP2 only at the shaft could also be seen. Further confirming that this heterogeneity may relate to the cell cycle, a similar staining variety (albeit no vesicle staining) was observed among ASC52telo cells, which are cycling as well, whereas, in hTERT-RPE1 cells, which were starved and thus, synchronized, the variation was not visible, instead only the stage with microtubules and cilia staining was observed (Figure 4G).

To explore this hypothesis, we trained a machine-learning model to predict the cell cycle stage by predicting nuclear GMNN expression for images of RPTEC/TERT1 cells (Figure 4H). Nuclear GMNN expression is low in G1, moderate in S, and high in G2 phase^88^. We validated our model by comparing GMNN predictions to stainings for other cell cycle markers, such as KIF18B (accumulating in the nucleus during S and towards G2), CDT1 (pronounced in G1, decreasing in S), and the cyclin CCNA2 (peaking in late G2) and observed relationships as assumed based on our previous cell cycle studies ^89^ (Figure 4H). Applying this model to the images acquired in our study for CKAP2 in RPTEC/TERT1 cells we observed that cells featuring cilia with CKAP2 localization were predicted to be in S / early G2 (Figure 4I). Cells with G1 predictions did not show CKAP2 in cilia. To confirm that our observation was not due to the limited number of cells analyzed, we repeated the experiment, acquiring and analyzing 150 3D images revealing 4,149 cells and confirming that ciliary CKAP2 was significantly enriched in cilia of S and G2 cells (Figure 4J).

In summary, compared to other organelles and cellular structures, the primary cilium displays remarkable heterogeneity in multiple regards, both in terms of cell type specificity and in terms of cell-to-cell heterogeneity and subciliary distribution. Our atlas provides the data to further delineate this heterogeneity and relationships of ciliary protein localization and cell states - as we demonstrate by exploring cell-cycle related ciliary CKAP2 protein localization.

### Ciliary subcompartments with unique cellular functions

Spatiotemporal regulation is not limited to cilium versus other cellular compartments, but occurs also within the cilium. The existence of subciliary microdomains for further signaling organization has been suggested ^1^. In our manual annotation scheme we distinguished four different types of ciliary domains: tip, cilium, transition zone, and basal body, for which a comparative GO Biological Process enrichment analysis contrasted against cell body regions revealed GO terms uniquely attributable to each subciliary regions (Figure S4). Yet, thanks to the high spatial image resolution in our data set, we could explore subciliary protein localization beyond this manual scheme through CiliaQ-based image analysis ^90^. We used CiliaQ to create protein localization profiles for each cilium. The profiles confirmed our annotation scheme and showed a high variance for each annotation class, especially when including multilocalizing proteins (Figure 5A). This variance results from the diverse subciliary localization patterns that we observed (Figure 2D), and from the heterogeneity on the single cell level as well as the multilocalization of proteins, both presented above for signaling proteins (Figure 3B) and enzymes (Figure 3E).

**Figure 5:**
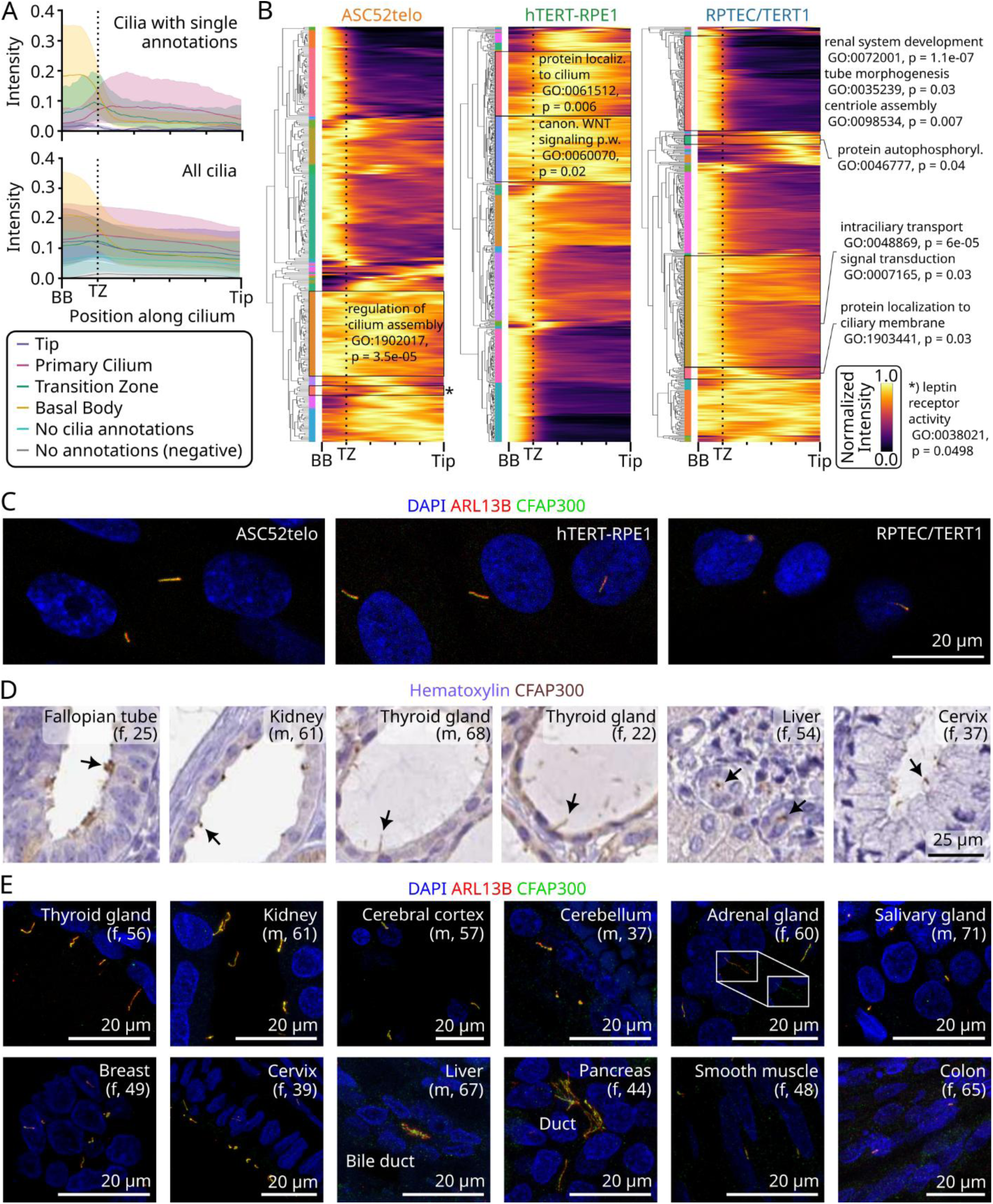
Ciliary protein domains and a new ciliary marker protein. **A)** Median (solid line) and 25% / 75% percentile (shadow) of normalized ciliary intensity profiles (scaled so that 0 (x-axis) represents the basal body (BB), 20 the transition zone (TZ), and 100 the ciliary tip (Tip), respectively, see STAR Methods). Top: Only ciliary intensity profiles from images annotated with only one of the four ciliary regions were included. Bottom: All analyzed cilia for approved stainings were included. “No cilia annotations” summarizes all profiles from images annotated as none of the ciliary regions but any of the other cellular regions. “No annotations (negative)” summarizes all profiles from images annotated as not showing staining. **B)** Hierarchical clustering of ciliary intensity profiles. Each row corresponds to the average ciliary intensity profile of one protein. Only proteins annotated as localizing to at least one of the four ciliary substructures in the respective cell line are included. Each averaged intensity profile was projected to a normalized intensity of 0 to 1 by dividing by the maximum intensity in the averaged profile. The x-axis represents the cilium from the center of the basal body (BB) to the center of the ciliary tip (as in A). TZ / dashed line indicates the transition zone. Intensity profiles were clustered individually for each cell line (see STAR Methods). The colored bars on the left indicate the proteins / profiles encapsulated within a cluster (colors are randomly assigned and do not match across cell lines). Each cluster containing at least five proteins / profiles was functionally investigated through gene ontology (GO) analysis. For all results, see Table S6; Selected clusters and GO terms are indicated in the figure. **C)** Example images for stainings of CFAP300, the only protein for which we observed staining only in the cilium. **D)** Example images from IHC stainings on human tissues (sex and age indicated in brackets) with the CFAP300 antibody (Image source: Human Protein Atlas). **E)** Confocal microscopy images (Maximum intensity projection of z-stack) from different human tissues (sex and age indicated in brackets) stained with a CFAP300 antibody, cilia (ARL13B antibody), and nuclei (DAPI). Inset shows the indicated region again with only the DAPI and CFAP300 channel displayed.

To unbiasedly explore subciliary protein localization patterns and scrutinize the existence of subciliary microdomains, we computed average ciliary localization profiles for each protein and cell line and hierarchically clustered these profiles (see Materials and Methods, Figure 5B, Table S6). In fact, localization patterns could be distinguished into at least 20 different kinds per cell line. The localization clusters align with distinct molecular and cellular functions of the proteins: For example, a cluster of ciliary intensity profiles in ASC52telo cells was related to “leptin receptor activity” (p=0.0499), a cluster in hTERT-RPE1 cells was enriched for proteins related to the canonical Wnt signaling pathway (p=0.02), and clusters in RPTEC/TERT1 cells related to renal system development (p=1.1e-07) and tube morphogenesis (p=0.03) (Figure 5B, Table S6). Profiles with high intensity at the cilium just above the transition zone were related to protein localization to the ciliary membrane (p=0.03) in RPTEC/TERT1 cells. And profiles enriched below the ciliary tip were enriched for proteins engaging in “protein autophosphorylation” (p=0.04) like TAOK1 (Figure 3B). Our analysis shows that proteins involved in similar molecular and cellular functions show characteristic ciliary localization patterns, supporting the existence of ciliary microdomains.

### A stable protein marker for cilia

Protein multilocalization and localization heterogeneity of ciliary proteins render it difficult to find suitable marker proteins for cilia imaging assays. A cilia marker protein that is stably and exclusively localizing along the whole primary cilium could greatly aid in identifying cilia in tissues and cells in research. Common cilia markers such as ARL13B, ADCY3, and acetylated tubulin come with the disadvantage of being signaling molecules and/or also localizing to other subcellular structures. ADCY3 antibodies appear to not label consistently cilia on all cell types ^85,91^, in line with our own observation of a high diversity of ADCY3 localization across the three studied cell lines (Figure 4B). ARL13B also has cytosolic roles ^92,93^, and, in particular, posttranslational tubulin modifications of the axonemal cytoskeleton, *i.e.*, acetylation, are known to be heterogeneous across cell types and also strongly label cytoskeletal structures in some cells, *i.e.*, neurons ^91,94^. In this study, only one single protein - besides the established marker ARL13B - was stably and exclusively identified in cilia, across all three cell lines: CFAP300 (Figure 5A). Mutations in *CFAP300* have been recently reported in patients with motile ciliopathy phenotypes and abnormal motile cilia structure, albeit the function of CFAP300 is not fully understood ^95–101^. We here demonstrate that CFAP300 is a suitable marker for primary cilia across multiple human tissues. For example, the antibody targeting CFAP300 shows a specific staining in immunohistochemical (IHC) DAB stainings in the fallopian tube (motile cilia), the kidney (primary cilia), the thyroid gland (primary cilia), the liver (primary cilia of cholangiocytes), or the cervix (primary cilia) (Figure 5B). Immunofluorescence (IF) co-staining with ARL13B and PCNT show that CFAP300 clearly marks primary cilia in the kidney, the thyroid gland, the cerebellum, the cortex, the adrenal gland, the salivary gland, the breast, the cervix, the liver (cholangiocyte cilia), the pancreas, the smooth muscle, and the colon, where mesenchymal cells are ciliated (Figure 5C). Of note, the CFAP300 antibody is highly validated, both, through specific binding to overexpressed tagged protein and through an external study that observed no staining when applying our HPA antibody to cells from a patient with a CFAP300 mutation causing a truncation of part of the protein, including the epitope region used to generate the antibody ^97^. We suggest CFAP300 as a new primary cilia marker, complementary to the existing commonly applied marker panel.

### Exploring new ciliary genes and their disease associations

To evaluate the functional significance of the newly identified ciliary proteins and to uncover potential clinically relevant ciliopathy genes, we compared the 715 proteins detected in primary cilium (including tip and transition zone) or basal bodies with phenotype annotations in the Online Mendelian Inheritance in Man (OMIM) database ^102^. This analysis resulted in three distinct groups: (1) Genes found in OMIM with a known phenotype (n=324), (2) Genes found in OMIM without an associated phenotype (n=338), and (3) Genes not listed in OMIM (n=53 (Table S7, Figure 6A). Most genes (217 of 324 in group 1, 333 of 338 in group 2, and 52 of 53 in group 3) were not labeled as ciliopathy-causing genes in OMIM (Figure 6A).

**Figure 6:**
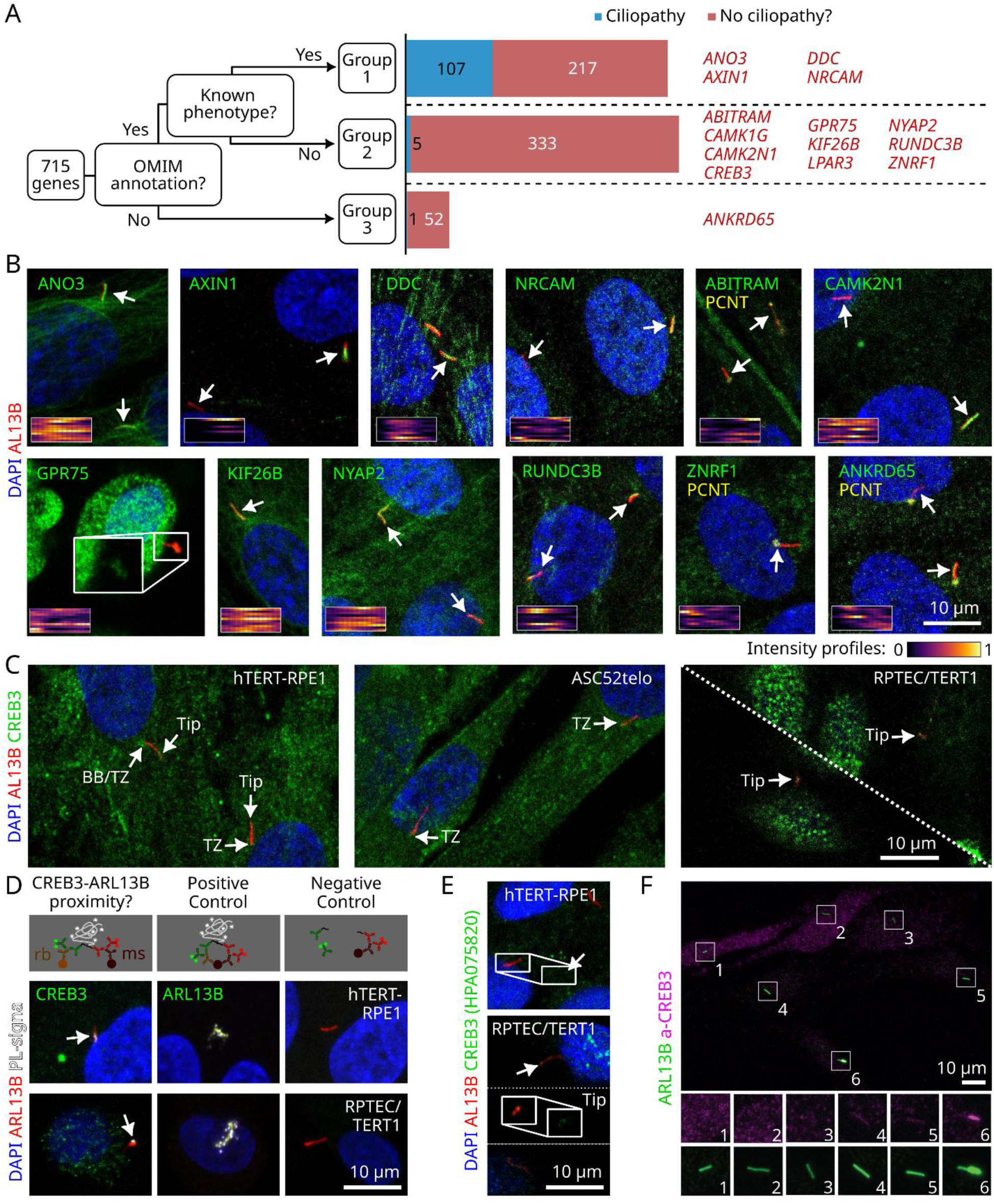
Potential clinical relevance of the newly revealed ciliary proteome. **A)** Flow chart for splitting the 654 genes into three groups based on their OMIM annotations. Additionally, the genes were classified by being established ciliopathy proteins or not. Example genes for each group are written on the right, and shown in B. **B)** Example images for staining the proteins for genes pointed out in A. Cell lines for images: Starved hTERT-RPE1 cells: NRCAM, DDC, NYAP2, CAMK2N1, ZNRF1, AXIN1, KIF26B, ANO3; RPTEC/TERT1 cells: GPR75; ASC52telo cells: ABITRAM. Insets show the intensity profiles of the protein of interest for 10 individual example cilia (each row represents one cilium) from the center of the basal body (left) to the tip (right), revealed by CiliaQ analysis (see STAR Methods). Magnified view for GPR75 shows only the green channel. **C)** Example images from stainings of CREB3 in hTERT-RPE1 cells (serum-starved), ASC52telo cells, or RPTEC/TERT1 cells. The white interrupted line distinguishes two images that were placed adjacently to show two cilia. Abbreviations: TZ = transition zone. BB = Basal Body. The basal body PCNT marker was not shown to better visualize the localizations of CREB3 - The PCNT marker was however used to correctly mark BB and TZ positions with arrows. **D)** Example images from a proximity ligation assay revealing that CREB3 and ARL13B localize within 40 nm proximity at cilia (see STAR Methods). **E)** Example images from stainings of CREB3 in hTERT-RPE1 cells (serum-starved) or RPTEC/TERT1 cells using an alternative antibody to C (HPA075820). **F)** Example image from a staining of hTERT-RPE1 cells overexpressing human CREB3 fused to an N-terminal alpha tag. An ARL13B antibody was used to label cilia and a fluorophore-conjugated alpha-tag binder to label the overexpressed CREB3. The cells showed varying levels of CREB3 in the cytoplasm and cilia.

Intersecting the list of genes with unknown phenotypes and the list of newly discovered cilia genes yielded a list of genes that inspire new perspectives on cilia, 14 of which we considered strong candidates to further explore in the future and thus highlight here (Figure 6A). Most of these showed a high spatiotemporal variation across cilia, and surprisingly, many of these proteins are enriched in the brain tissue or fulfill key roles in neurons. These include: the cell adhesion molecule NRCAM (Figure 6B), which is involved in cell-cell contacts and axon outgrowth in the peripheral- and central nerve system ^103^; the aromatic-L-amino-acid decarboxylase DDC (Figure 6B) involved in neurotransmitter metabolism; the Neuronal tyrosine-phosphorylated phosphoinositide-3-kinase adapter 2 NYAP2 (Figure 6B), which belongs to a family of proteins that regulates neuronal morphogenesis ^104^; The calcium/calmodulin-dependent kinase CAMK1G (Figure 3B), which can act in a calcium/calmodulin dependent signaling cascade important for axon formation ^105^; the calcium/calmodulin dependent protein kinase II inhibitor 1 CAMK2N1 (Figure 6B), which inhibits CAMK2, a key kinase in many neuronal functions ^106^. Furthermore, we identified the E3 ubiquitin ligase ZNRF1 at the transition zone (Figure 6B). ZNRF1 is enriched in the nervous system and targets various receptors ^107^, but also interacts with tubulin ^108^. The little studied actin binding transcription modulator ABITRAM was observed in individual cilia of ASC52telo and hTERT-RPE1 (Figure 6B).

We found the GPCR LPAR3 in individual cilia, as described above (Figure 4F). Interestingly, LPA signaling has recently gained more attention in ciliogenesis regulation and glioblastoma, yet focusing on LPA detection through LPAR1 but lacking clarity on where endogenous LPARs localize ^109–111^. Furthermore, in a study using zebrafish as a model organism, LPAR3 had been recently implicated to be crucial for establishing Left-Right asymmetry ^112^, where cilia are known to play important roles, and brain functions ^113^. Exogenous FLAG-tagged LPAR1 and LPAR3, introduced by overexpression, have been shown to localize to cilia of immortalized astrocytes ^110^. We now reveal endogenous LPAR3 in human cilia, supporting the relevance for LPA signaling in cilia. Another highly interesting GPCR that we identify in cilia is GPR75 (Figure 6B). GPR75 has recently gained a lot of attention due to human GPR75 variants protecting from obesity ^114^. In mice, tagged GPR75 had been confirmed in cilia in hypothalamic neurons and linked to food intake ^115^. We here confirm endogenous human GPR75 in cilia. Since ciliary GPR75 detection is in the low SNR range, we further confirmed ciliary localization with a highly sensitive proximity-labeling assay, which identified GPR75 and ARL13B within 40 nm proximity at the tip of cilia (Figure S5A).

An additional protein that we revealed in cilia is AXIN1 (Figure 6B), reported to be part of the beta-catenin destruction complex and to regulate beta-catenin levels and modulate Wnt signaling ^116–118^. Just recently, Wnt signaling has been highlighted as an additional signaling pathway in primary cilia ^119^. We add AXIN1 to the picture of the Wnt signaling pathway in cilia. Interestingly, those cilia showing AXIN1 staining in hTERT-RPE1 cells were off focus compared to the cell nucleus, sticking into the liquid space of the culture well (Figure 6B). Furthermore, we revealed in cilia the kinesin KIF26B (Figure 6B), the calcium-dependent regulator of potassium channels ANO3 (Figure 6B), and the uncharacterized proteins RUNDC3B (Figure 6B), and ANKRD65 (Figure 6B).

To further validate the presence of the newly identified ciliary proteins lacking validation through other literature, studies, or described ciliopathy-related phenotypes, we attempted validation with secondary antibodies targeting alternative antigenic sequences or through recombinant protein expression of tagged versions of the candidate protein. This allowed us to validate ciliary localization for ABITRAM (Figure S5B), ANO3 (Figure S5C), NRCAM (Figure S5D), HTR7 (Figure S5E), BMP2 (Figure S5F), and CAMK1G (Figure S5G,H).

Lastly, we investigated genes from groups two and three (Figure 6A) in clinical samples analyzed by whole genome sequencing (WGS) ^120,121^. Clinical WGS data from three patient groups were analyzed: 1) patients with suspected ciliopathy without a causative variant found (n=157), 2) fetuses with an ultrasound-detected malformation without a causative variant found (n= 470) and 3) patients who had undergone a clinical patient-parental trio WGS analysis due to a suspicion of a potential syndrome but without a causative variant found (n= over 1,500). Patients with different symptoms, not only symptoms fitting a ciliopathy disorder, were included in group 3. We searched for potential clinically relevant single nucleotide variants in the 354 genes in groups two and three (Figure 6A, Table S7) in the WGS data from these three patient groups by one gene at a time, filtering for frameshift variants, nonsense variants and predicted splicing variants in homozygous, compound heterozygous or *de novo* states. Interestingly, we identified one individual in group 3 for whom a *de novo* two-nucleotide deletion was detected in *CREB3* (NM_006368:c.810_811del) causing a premature stop codon (p.Ser271*) (Comprehensive clinical description in Note S1). The individual was a 2-year-old boy with a syndromic clinical presentation including optical nerve hypoplasia (ONH), a small pituitary gland, retinal dystrophy, growth failure, skeletal abnormalities, and endocrine dysfunction (growth hormone deficiency and hypothyroidism). His other symptoms include retinal changes, nystagmus, low bone density, short stature, and hypoglycemia. We observed CREB3 in the cilia of RPTEC/TERT1 cells with an enrichment at the ciliary tip, at the transition zone region in ASC52telo cells, and at basal bodies or the transition zone, or at the tip in starved hTERT-RPE1 cells (Figure 6C). Since this localization pattern is difficult to identify in cells with protein localization to other cellular compartments, we validated CREB3 localization to the cilium using multiple approaches. First, we applied a proximity-ligation-based assay confirming that ARL13B and CREB3 co-localize in cilia within a distance of 40 nm (Figure 6D). Second, we confirmed the localization of CREB3 to the cilium using an independent (non-overlapping epitope) antibody (Figure 6E) and through overexpression of tagged human CREB3 (Figure 6F). Our findings, combined with the patient’s phenotype overlapping with features of ciliopathies, suggest CREB3 as a new candidate ciliopathy-associated gene.

## Discussion

Our study provides the first insight into single-cilia proteomic heterogeneity and a unique resource for the cilia community to develop new hypotheses about the cellular functions governed by the primary cilium and their underlying mechanisms. We resolve domains of heterogeneity, such as within and across cell populations, highlight the specificity of ciliary signaling pathways, reveal ciliary microdomains, identify a new ciliary marker, and employ our atlas to search for new ciliopathy candidate genes.

Systematic MS-based proteomics approaches have advanced our understanding of the ciliary proteome in selected cell types ^20–25^. However, such methods have been lacking the resolution to capture the variability of individual cilia (see introduction). In turn, our understanding of cell-type-specific ciliary proteins remained limited to a few proteins. For example, many years of mechanistic research, using genetic tools, cellular models, and model organisms have established that GPCRs localize to cilia with cell-type specificity ^122^. With our method we greatly advance the understanding of cilia as we show that a broad variety of ciliary proteins, not only receptors but also kinases like PI3K, AKT kinases, or calmodulin-dependent kinases, cytoskeleton-related proteins like Septins, and enzymes are in fact cell-type- and even single-cell-specifically localized to the cilium. The presence of downstream signaling machinery in cilia, such as kinases, stresses that the cilium is not just a cellular antenna but also a computer that integrates signals to compute a response decision, to be relayed to the cell body. It has been established for sperm cells that the local signaling machinery in the flagellum modifies the flagellar beating locally for steering the cell ^123^ - our data propose that this concept holds true for cilia as well.

We observed that 99% of the investigated ciliary proteins multilocalize. Since, when the ciliary proteins were not detected in cilia, they were still detected in other cellular structures, we suggest that ciliary specificity across cell types may be substantially driven by protein trafficking, in addition to gene expression. Ciliary import and export have been identified as a signal transduction mechanism, such as in Hedgehog signaling ^1^. Our data supports extending this concept to many more signaling pathways in cilia, as the related proteins heterogeneously localize to cilia in our assay. We propose that cells not only use ciliary import and export to transduce signals but to also rapidly fine-tune their antenna to individually sense their environment depending on their specific task in a cellular consortium. Our dataset serves as a starting point for further studying this hypothesis, e.g., through relating ciliary protein localization and cellular states, as we have demonstrated for the CKAP2-localization and its relation to the cell cycle.

Structurally, we go beyond the resolution of previous studies: we identify the distinct protein compositions at the tip, cilium, transition zone, and basal body in the same cell. GO enrichment analysis suggests that these regions are specialized for specific signaling functions, which may be modes of shaping the computational architecture of the cilium. Our analysis of protein localization pattern similarity along the cilium hints at spatial organization of ciliary proteins into functional microdomains, which may even be dynamic. This data set can serve as a starting point for informing systematic super-resolved cilia studies, for example using expansion microscopy or cryo-CLEM.

Importantly, the ciliary compartment is not just organized on the level of proteins. Beyond signaling molecules, we also reveal enzymes in cilia that may close gaps in our understanding of how the ciliary compartment is organized. For example, we reveal an enzyme involved in sterol synthesis, supporting the hypothesis that cilia locally metabolize their unique lipid environment for customizing or conducting signaling pathways ^77^. The presence of the other enzymes suggests that the cilium has the capabilities to locally modify and possibly metabolize many more molecules and proteins than yet known, which may be crucial to further study to understand their contribution to ciliary signaling pathways and functions. The heterogeneity of enzyme localization to cilia highlights the importance of single-cell spatially-resolved methods for protein, lipid, and metabolite measurements to further delineate the complex architecture of the ciliary compartment.

Beyond highlighting the cilium as a sensory signaling device, which features not only reception but also signal amplification and processing capabilities, our data also highlight other ciliary functions to be further explored. Interestingly, we detected the fewest cell-type-unique proteins in ASC52telo cells, with only 19 or 16 cell-type unique primary cilia or basal body protein localizations - much less than hTERT-RPE1 (83 or 95, respectively) or RPTEC/TERT1 cells (88 or 70, respectively). This might be due to technical limitations, as ASC52telo cells have shorter flat-lying cilia compared to the other two cell types and a significantly lower ciliation rate (Figure 1D). Despite this, the fraction of proteins not previously described for cilia in our candidate list was collected with a strong focus on signaling cascades. Based on our gene ontology analysis of ASC52telo cells and the fact that they have cilia only for a very short time due to their fast cell cycle, we hypothesize that the cilium of ASC52telo cells is less tuned to receptor-based signaling but possibly rather crucial for other functions, such as driving stemness and asymmetric cell division. Asymmetry may be produced by the ASC52telo cells retaining a ciliary vesicle or whole cilium during cell division. Thus, our data encourage us to next direct our antibody-based spatial proteomics methods to stem cells with an expanded candidate gene list tailored to stem cell functions. Furthermore, we surprisingly identified many neuron- and brain-related proteins in cilia, which put neuronal cilia into focus even though we did not study neuronal cells. Some of these proteins relate to synapses or axons, supporting the recent observations of the cilium being involved in synaptic transmission ^67^ and that the cilium is not just an organelle that is a free-floating antenna but that can also establish contacts with other structures, *i.e.*, synapses, for even more local signal reception.

Our study opens up many new avenues for ciliary research. First, we have tested and validated antibodies for 715 proteins in primary cilia in human cells. Most of these antibodies are commercially available and allow the field to probe ciliary signaling and structure in detail, for example, when investigating patient-derived cells to explore disease mechanisms. Second, we also show that our screen has the potential to be translated to deep studies of human tissues, which we pursued for revealing CFAP300 as a new ciliary marker. Notably, many of the other antibodies that we tested are also suitable for tissues, as they are already validated in IHC workflows in the Human Protein Atlas ^27,35^. Recently, we also showed that the antibody for ATF3, which we identified at the Basal Body, is also applicable to study the role of cilia in the developing human heart ^124^. Third, we generate a dataset that can provide new avenues for clinics to reveal potential new ciliopathies and better understand ciliopathies by showing where cilium-related proteins localize. As ciliopathies are rare and heterogeneous disorders, patients often remain undiagnosed for years, and the disease ontogeny remains unresolved since the disease-causing genetic variants are not located in known ciliopathy genes. Using our new atlas, we identified *CREB3* as a novel candidate gene in a young boy with blindness, severe short stature, and additional syndromic features. While this finding highlights *CREB3* as a promising candidate gene, no causality between the *CREB3*-variant and the patient’s symptoms can be established without further experimental studies. *CREB3*-related phenotypes from animal models ^125–127^, including somatotroph hypoplasia, dwarfism, as well as altered hypothalamic-pituitary stress pathways, align with the patient’s brain abnormalities and growth failure. This study involved a targeted search for candidate variants in a curated list of potential new cilia genes in selected clinical genomes as a proof-of-concept pilot. In the future, this approach could be extended to broader gene lists and larger patient cohorts to better characterize the spectrum of ciliopathy symptoms and the full range of potential causal genes.

Beyond *CREB3* and classical ciliopathies, our study opens up avenues for clinical cilia research. First, we confirm endogenous GPR75 in cilia, a recently discovered candidate for cilium-regulated satiety regulation ^114,115^. Second, we confirm several proteins encoded by genes showing genetic variants in autism patients to localize to human cilia ^128^. Third, our discovery of the new ciliary marker protein *CFAP300* has implications for clinical research. So far, genetic mutations in *CFAP300* have been linked to motile ciliopathy phenotypes in patients worldwide, CFAP300 has been highlighted as a highly conserved protein across ciliated organisms, functionally been attributed roles in dynein-arm assembly for motile cilia, and been described to show patterns resembling IFT-transport ^95–101^. We here show that CFAP300 localizes to primary cilia as well, and observe a spotty pattern on cilia that is often seen for IFT particles or cargos, in line with previous studies ^97^. Based on our data, we hypothesize that CFAP300 fulfills a general role beyond motile cilia, also in primary cilia. It remains puzzling that CFAP300 patients have so far been diagnosed only with motile ciliopathy symptoms. Noteworthy, multiple transcripts and protein isoforms of CFAP300 exist. It can be speculated that the different isoforms feature different functions and patient mutations affect not all isoforms or protein functions equally; or that CFAP300 mutations affecting the function in primary cilia are either lethal early in life or less crucial for life leading to non-diagnosis of minor primary cilia related phenotypes. Future functional studies will be required to delineate the functions of CFAP300 in different cilia. Nonetheless, this observation - together with our observation of motile cilia proteins like IQCA1 or HYDIN in kidney cilia - highlights that the distinction between motile and primary cilia is not as clear-cut as anticipated and that more attention could be directed to the overlap of primary and motile cilia functions and mixed clinical phenotypes.

In summary, we highlight the primary cilium as a cellular compartment with variable proteome and function. The ciliary compartment is likely regulated to bring together certain signaling molecules in time and space. We hypothesize that cells employ their cilia to customize sensation and signal transduction to individual cellular roles. Thereby, the primary cilium might be a space in which the cell can both tailor their environmental sensing abilities and customize the internal computation to allow precise cell type and state-tuned signaling. Our data suggest that there are multiple domains of ciliary heterogeneity potentially relating to cell states, such as cell cycle, ciliogenesis, signaling, and metabolic activity. Further deep analysis of our data set followed up by functional assays and live measurements of heterogeneous protein localization will allow the cilia community to further define these axes of heterogeneity. Hence, this study serves as a starting point for cilia systems biology, providing a template for modeling the cilium as an information processing unit. With all our data being publicly explorable at https://www.proteinatlas.org we pave new avenues for research on decoding the fundamental biology of human primary cilia and clinical implications such as in ciliopathies.

### Limitations of the Study

This study reveals ciliary heterogeneity based on three immortalized cell lines from different tissues. To what extent the heterogeneity translates to tissues remains to be further explored.

The study uses HPA antibodies, which have been designed to target unique pan-isoform regions of each protein, affinity purified, tested for off-target binding in affinity protein arrays, and extensively studied and validated in different paradigms, including stainings of tissues and cell types relatable to RNASeq expression data, Western Blots. Some antibodies were also further scrutinized through siRNA, protein-tagging, and MS-based validation studies. All stainings in this study underwent a multi-step evaluation process, in which the stainings are assessed by multiple skilled cell biologists for signs of unspecific binding, alignment with literature and other protein knowledge resources, and other staining data. Ultimately, annotated subcellular localizations were scored for their reliability as determined by skilled cell biologists based on available validation and literature data - the scores are provided on the Human Protein Atlas webpage. Table S8 summarizes part of the antibody validation data, sourced from HPA version v24. Since all validation data and the antigenic sequence used for immunization is publicly available on the Human Protein Atlas webpage, we encourage researchers interested in specific proteins to inspect the available validation data for drawing individual conclusions for their specific research case.

Noteworthy, the HPA is a long term project, in which the antibodies and stainings, including those in this study, can be reevaluated if new data emerge. Additionally, more proteins and antibody stainings may be added to the HPA cilia section in the future.

We caution that proteins not identified in cilia in this study may still be present in cilia. Our approach was targeted, and, due to feasibility, limited to 1,992 candidate proteins. The selection of candidate proteins relied on the availability of antibodies - we may miss data on ciliary proteins for which validated antibodies were lacking. Furthermore, our screen suggests that 645 proteins from the candidate list did not localize to cilia in these cell lines based on validated antibodies. However, other reasons for not seeing them in cilia may be extremely low abundance, signaling-dependent or cell-type-specific cilia localization, post-translational modifications interfering with antibody binding, or limited accessibility of the antigenic sequence in cilia. We conclude that absence of a protein in our study or atlas does not necessarily indicate that it is not present in cilia.

Due to the screening nature and the extensive efforts for data generation, this study relies on single-shot stainings. By automatizing the staining and imaging pipeline, human error is minimized. The human-based evaluation pipeline includes assessment for potential staining errors or artefacts, resulting in data exclusion and staining repetition.

## Resource availability

### Lead contact

Further inquiries should be directed to the Lead Contact, Emma Lundberg.

### Materials availability

The antibodies generated within the HPA are commercially available, purchasable through their HPA ID (e.g., HPA038585 for the antibody targeting CFAP300) as catalogue number. Cell lines, other antibodies and reagents used in the study are commercially available and product information can be found in the methods section.

### Data and code availability

The data presented in this study, including images and annotation, is available in the current version of the Human Protein Atlas (v24) or will be made available in the upcoming version of the Human Protein Atlas (v25) at https://www.proteinatlas.org.

All software and code used in this study is available on GitHub (https://github.com/CellProfiling/HPA_Cilia_Study_Code) unless otherwise stated in the methods part.

## Acknowledgements

This research was funded by grants from Erling Persson Foundation, Göran Gustafsson Foundation, Chan-Zuckerberg Biohub, and Knut and Alice Wallenberg Foundation to EL (KAW 2021.0346) as well as to MU (HPA). JNH was supported by a Postdoctoral Fellowship from the Wenner-Gren Foundations, and by an EMBO Postdoctoral Fellowship (ALTF 556-2022). We thank the Lavis lab and Open Chemistry team (Janelia) for the gift of JF and JFX dyes.

## Author contributions

JNH, UA, and EL conceived the project and designed the study. JNH, UA, AJ, CV, KT, AMC, EP, AM, JF, and AB conducted experiments. UA, JNH, KT, JF, EL, and DM annotated images. JNH, UA, EL, HS, KK, EW, TL, CW, SHS, and DM analyzed and interpreted data. MU and CLB contributed to the discussion and critical reagents. JNH, UA, and EL wrote the paper. ADV and AL supervised the clinical analysis. JNH and KK visualized the results and prepared figures. EL supervised the project. KVF, FJ, and UA built the cilia atlas in HPA. All authors edited and commented on the manuscript.

## STAR methods

### Key resource table

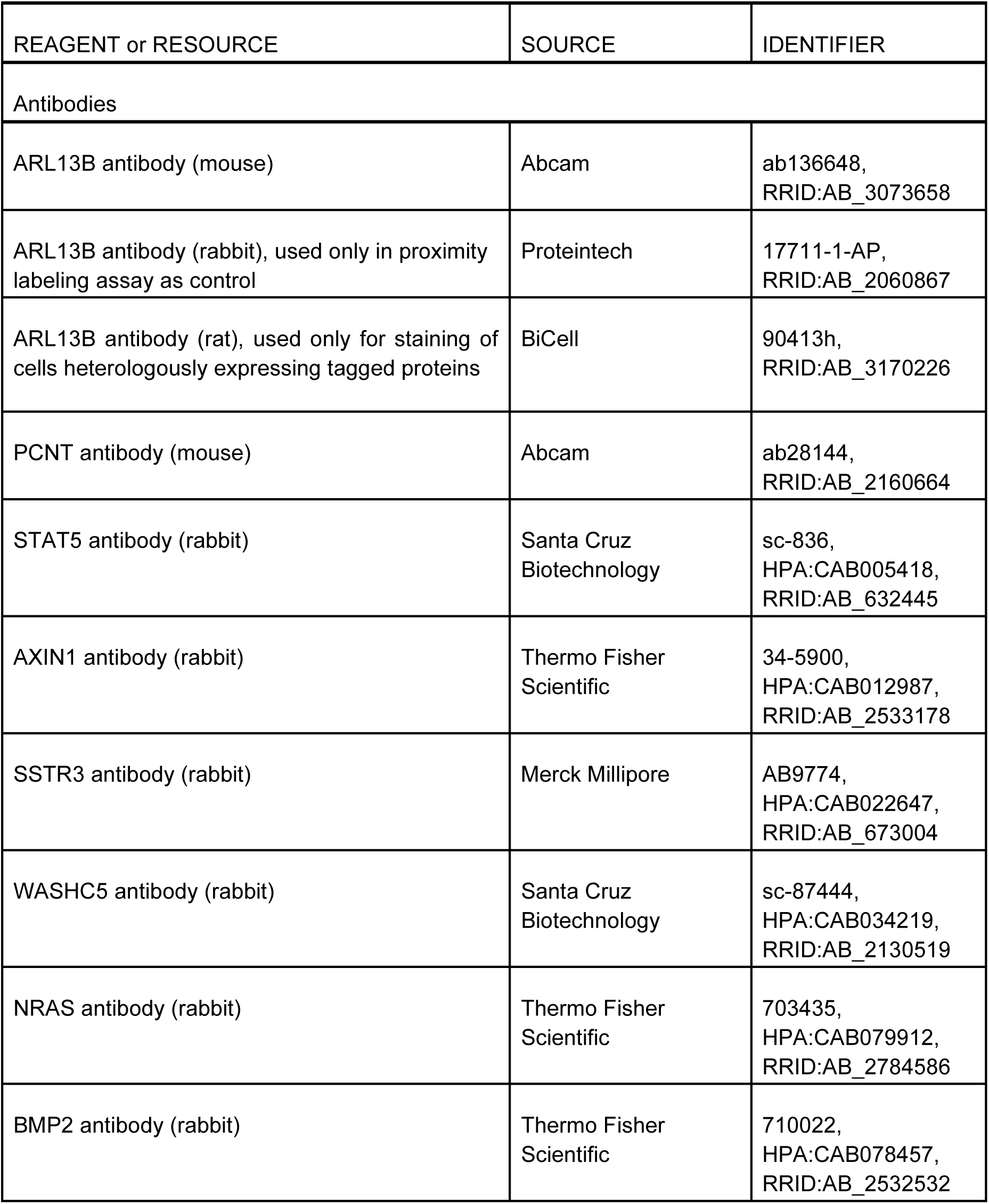
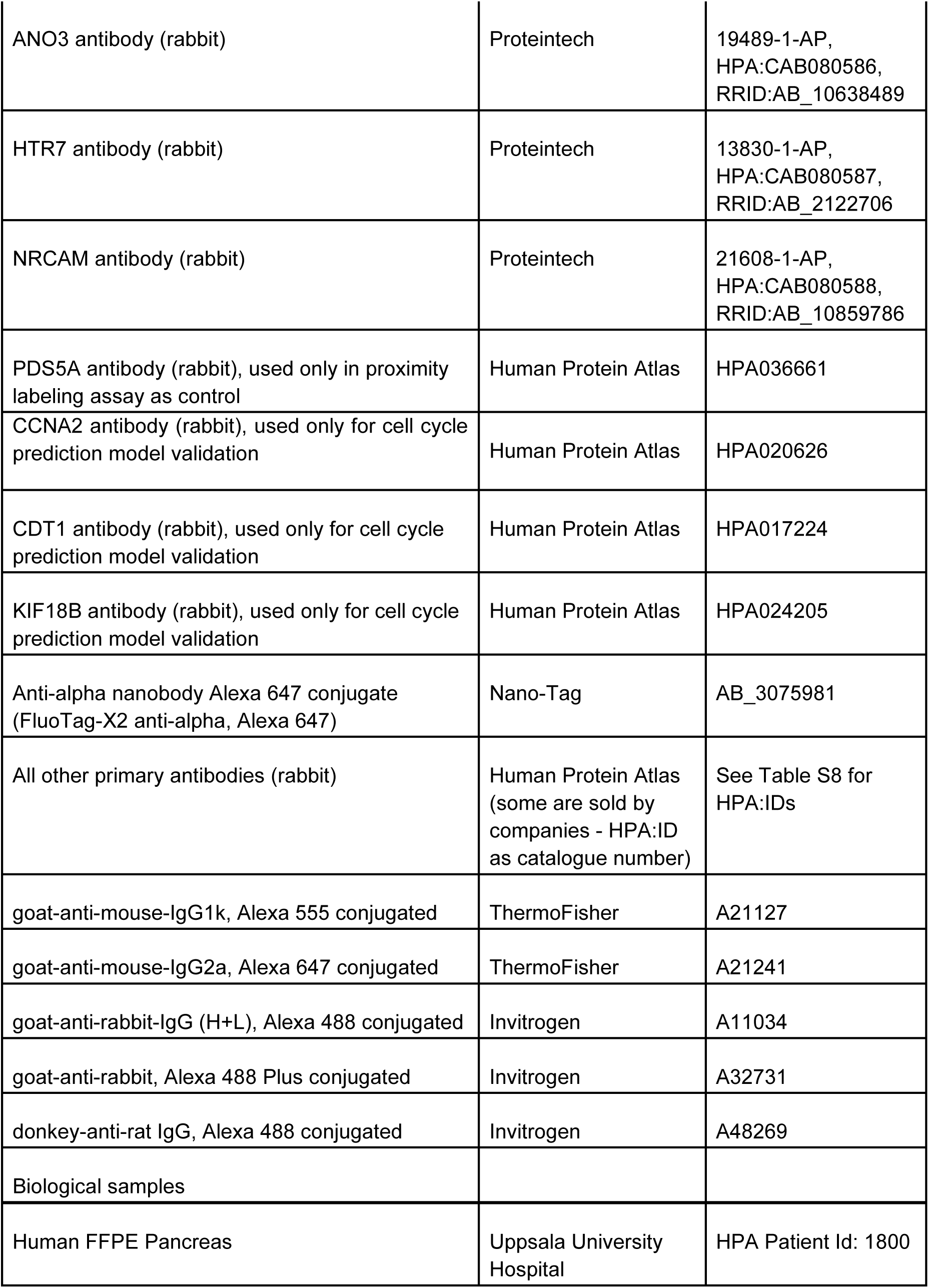
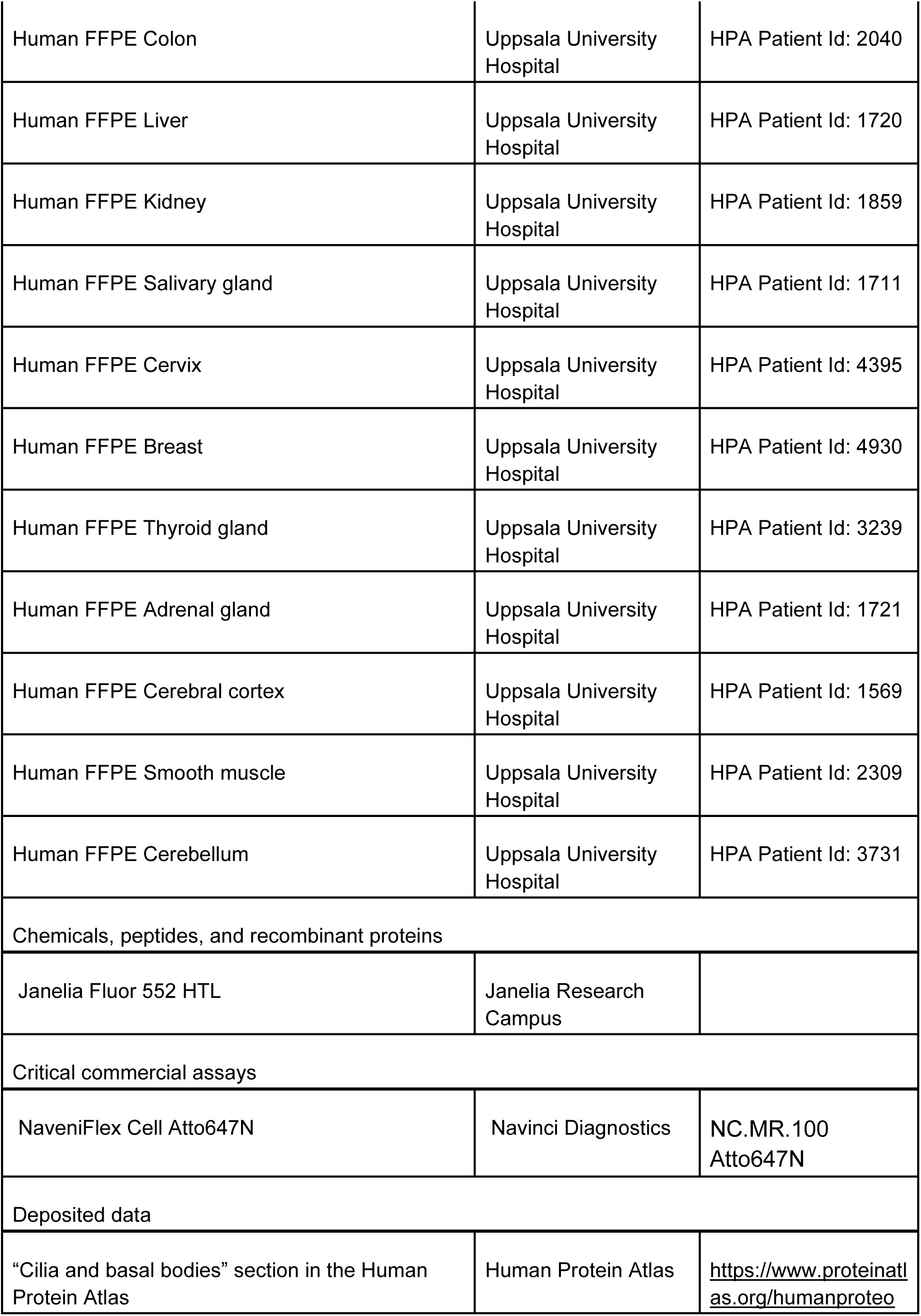
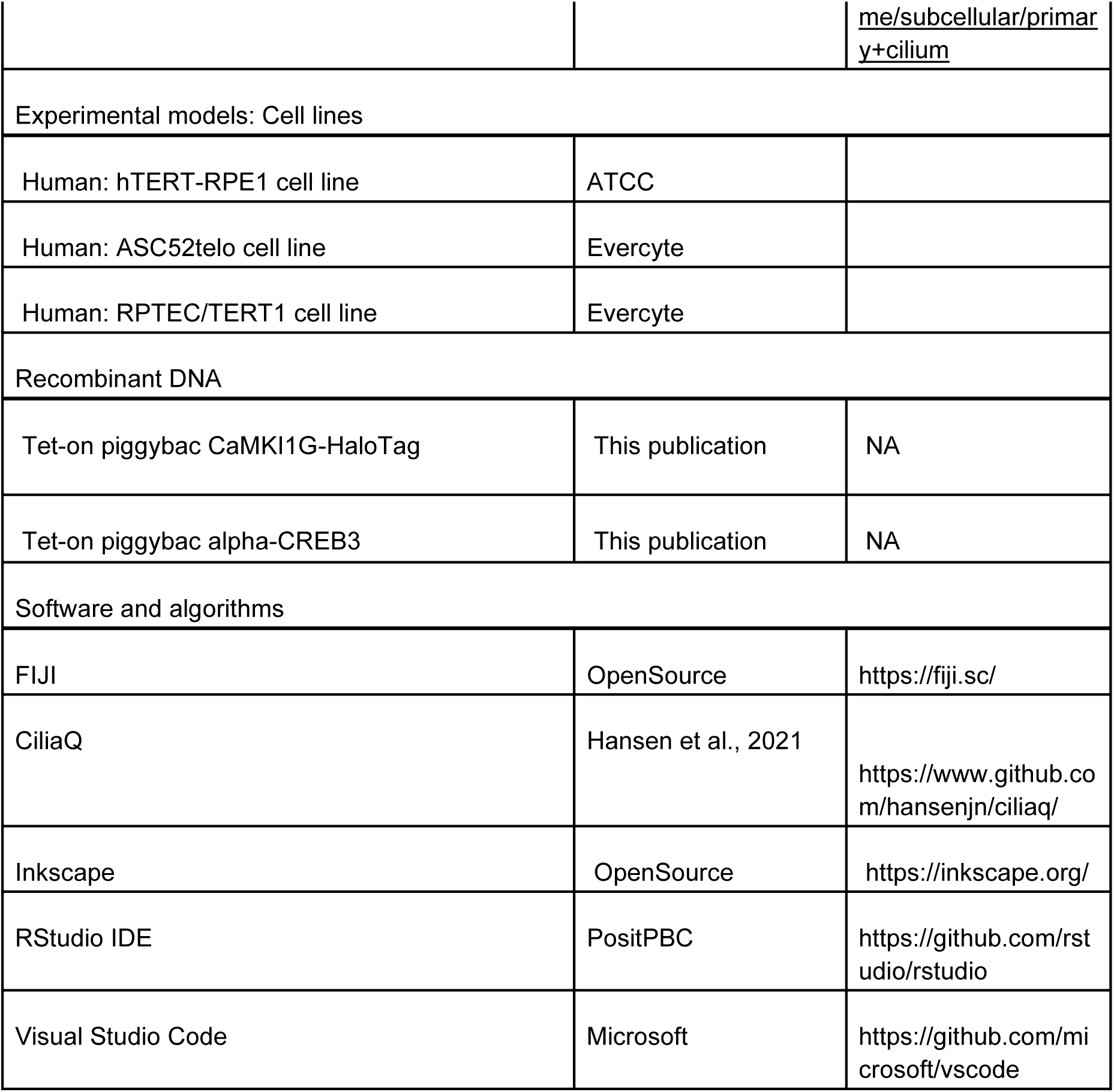

### Experimental model and study participant details

#### Cell culture

hTERT-RPE1 cells were acquired from ATCC and cultivated in DMEM/F12 (Thermo Fisher Scientific Inc, GibcoTM, Cat. no. 11320033) + 10% FBS (Merck KGaA, Sigma-Aldrich, Cat. no. F7524) at 37°C and 5% CO2. ASC52telo and RPTEC/TERT1 cells were acquired from Evercyte GmbH (Vienna, Austria). ASC52telo cells were cultivated in Endothelial Cell Growth Medium-2 (Lonza, Cat. no. CC-3162) supplemented with 4% FBS (Merck KGaA, Sigma-Aldrich, Cat. no. F7524) and 200 ug/ml G418 (G-418 Solution, Merck KGaA, Cat. no. 04727878001). RPTEC/TERT1 cells were cultivated in ProxUp2 medium (Evercyte, MHT-003-2).

#### Human tissue samples

The tissue samples were processed following Swedish laws and regulation, obtained from the Department of Pathology, Uppsala University Hospital, Sweden, as included in the sample collection governed by the Uppsala Biobank (http://www.uppsalabiobank.uu.se/en/). Informed consent was obtained from all individuals. All human tissue samples used were anonymized in accordance with an approval and advisory report from the Uppsala Ethical Review Board (Dnr Ups 02-577). All samples were obtained from archival tissues from adult human subjects, for which the ethical permit only allowed access to age, gender and histopathological information of the original tissue sample. Only histologically normal regions were included in the present investigation. Samples were used for generation of tissue microarrays (TMAs), including three individuals for each tissue type. When applicable, both male and female subjects were used.

### Experimental details

#### Selecting ciliary candidate proteins

Lists of genes or proteins detected in primary cilia, motile cilia, and/or flagellum in humans and across other species were extracted from databases in the format of supplementary tables, PDF-files, or exportable csv files. The data from ChlamyFP.org (Human) were downloaded on July 10, 2024. The files were next imported into R and merged into a master list (Table S1) by conversion of the protein or gene identifiers into Ensembl gene IDs using the R package biomaRt (version 2.56.1) ^129^. Conversions failing in bioMart were converted - if possible - by automated queries to the Ensembl REST API (https://rest.ensembl.org). The data overlap was plotted using the library Plotrix (version 3.8.4). To reduce the list to the candidate list of 1,992 genes, we selected targets based on (1) whether the gene is expressed in the cell lines we aimed to study (using HPA internal cell line RNAsequencing data), (2) whether they represent gold standard cilia proteins, (3) whether the HPA antibody library provided a good, validated antibody for the protein of interest, and (4) the discovery potential of the protein of interest, with a strong focus on revealing signaling proteins in cilia. For completeness of our characterization of the ciliary proteome, we included stainings of nine proteins, performed using the same pipeline, but for a project focused on cilia in autism^128^, into data analysis. Specifically, these are 3D images from stainings for ADNP (Antibody: HPA050560), CHD8 (HPA076133), DLG4 (HPA071006), GIGYF1 (HPA023995), GRIN2B (HPA052160), SATB2 (HPA029543), STXBP1 (HPA008209), SYNGAP1 (HPA038374), XPO1 (HPA042933).

#### Staining prioritization and experiment order

We started by staining the one or two cell lines with highest RNA expression (HPA internal cell line RNA expression database) and did not pursue further staining in the other cell line(s) if the staining / antibody was deemed unspecific or unreliable or did not show cilium or basal body localization of the protein and in the other cell lines protein expression was not detected on RNA level. This resulted in not all proteins being stained in all three cell lines. Considering only finally approved stainings, 737 genes were stained for in all three cell lines, 336 genes were stained for in two of three cell lines, and 287 genes were stained for in only one cell line. For all conclusions drawn on inter-cell line diversity, only genes that have been stained in all three cell lines were included.

#### Immunocytochemistry

Cells were seeded in 80 µl / well of the respective growth medium on 96-well glass-bottom plates (Greiner Sensoplate Plus, Greiner Bio-One, Germany) coated with 50 μl of 12.5 μg/ml human fibronectin (Sigma Aldrich, Darmstadt, Germany) and cultivated at 37 °C and 5% CO2. For hTERT-RPE1 and ASC52telo cells, approximately 10,000 cells/well were seeded. For RPTEC/TERT1 cells, approximately 30,000 cells/well were seeded. hTRET-RPE1 cells were starved 24 hours after seeding by medium exchange to DMEM/F12 + 0.2% FBS, and fixed 48 hours after start of starvation. RPTEC/TERT1 cells were fixed 24 hours after seeding. ASC52telo cells were fixed 48 hours after seeding. For fixation, the cells were washed with phosphate-buffered saline (PBS, pH 7) and fixed with 40 μl 4% ice-cold paraformaldehyde (PFA: Sigma Aldrich, Darmstadt, Germany) dissolved in PBS for 15 min. Cells were washed with PBS and permeabilized with 40 μl 0.5% Triton X-100 (Sigma Aldrich) in PBS for 3 × 5 min. Rabbit polyclonal HPA antibodies targeting the proteins of interest were dissolved to 2–4 μg/ml in blocking buffer (PBS + 4% FBS) containing the cilia marker antibodies against ARL13B (Abcam, ab136648, mouse IgG2a, 1:2000) and PCNT (Abcam, ab28144, mouse IgG1, 1:2000). After washing with PBS, the diluted primary antibodies were added (40 μl per well), and the plates were incubated overnight at 4 °C. After overnight incubation, wells were washed with PBS for 3 × 10 min. Secondary antibodies, goat-anti-mouse-IgG1k-Cross-Adsorbed-Alexa555 (ThermoFisher, A-21127, 1:2000), goat-anti-mouse-IgG2a-Cross-Adsorbed-Alexa647 (ThermoFisher, A-21241, 1:1000), and goat-anti-rabbit-IgG (H+L)-Highly-Cross-Adsorbed-Alexa488 (Invitrogen, A11034, 1:800) diluted in blocking buffer were added and the plates were incubated for 90 min at room temperature. Next, cells were incubated with 572 nM = 200 ng/ml of DAPI (4’, 6-diamidino-2-phenylindole) in PBS for 10 min, and washed 3 x 5 min with PBS.

#### Antibodies

The antibodies used in this study were generated within the HPA project except for the marker antibodies for cilia (targeting ARL13B, Abcam, ab136648, RRID:AB_3073658) and basal bodies (targeting PCNT, Abcam, ab28144, RRID:AB_2160664) and antibodies for STAT5 (HPA internal ID: CAB005418, Santa Cruz Biotechnology, sc-836, RRID:AB_632445), AXIN1 (HPA internal ID: CAB012987, Thermo Fisher Scientific, 34-5900, RRID:AB_2533178), SSTR3 (HPA internal ID: CAB022647, Merck Millipore, AB9774, RRID:AB_673004), WASHC5 (HPA internal ID: CAB034219, Santa Cruz Biotechnology, sc-87444, RRID:AB_2130519), and NRAS (HPA internal ID: CAB079912,, Thermo Fisher Scientific, 703435, RRID:AB_2784586). Table S8 lists the HPA antibodies applied in the screening approach, the gene symbols and Ensembl IDs for the proteins they bind, dilution factors applied, and a summary of validation data (as present in version v24 of the HPA). Some antibodies are able to bind multiple similar proteins, as specified in the table. All HPA antibodies are rabbit polyclonal antibodies generated using recombinant protein epitope signature tags (PrEST) as antigens for immunization ^130^. The PrESTs are designed to enable the antibody to bind specifically to as many isoforms of the target protein as possible. The resulting antibodies were affinity purified using the PrEST antigen as an affinity ligand. Commercial antibodies were provided by the suppliers and used according to the supplier’s recommendations.

As previously described, the antibodies were initially quality controlled for sensitivity and lack of cross-reactivity to other proteins using western blotting and protein arrays ^130^. Antibodies that passed the initial quality assessment, with no literature contradicting the results obtained in immunofluorescence (IF), were labeled ‘approved’. If there was independent data in UniProt supporting the results in IF, the antibody was labeled ‘supported’. For enhanced application-specific validation, we used the strategies outlined by the International Working Group for Antibody Validation (IWGAV) ^131^, including i) validation by gene silencing/knockout, ii) validation by co-staining of fluorescent protein-tagged protein expressed at endogenous levels, iii) validation by independent antibody targeting a different epitope, and iv) capture validation by mass spectrometry ^131,132^. If the antibody had been validated according to the IWGAV guidelines, it was labeled ‘enhanced’. A comprehensive summary of all antibody validation data for each gene is available through the Human Protein Atlas webpage from the “Antibody and Validation” tab provided on each gene page or through a link constructed as follows: https://www.proteinatlas.org/ <INSERT ENSEMBL GENE ID here>/summary/antibody.

#### Fluorescence image acquisition of cells

All fluorescent images of cells were acquired with an inverted Leica SP8 confocal microscope equipped with a motorized and piezo z stage, an HC PL APO CS2 63×/1.20 water-immersion objective (Leica Microsystems, Mannheim, Germany), Laser Diodes 405, 488, 552, and 638, and HyD detectors. The settings for each image were Pinhole 1 Airy unit, 16-bit acquisition, 1.28x zoom, 2048x2048 pixels field of view, 700 Hz scan speed, resulting in a pixel size of 0.07 μm. All channels were detected on HyD detectors in multiple individual series to reduce bleed through: (1) DAPI and Cilia (Alexa 647) were detected, (2) the protein of interest was detected (Alexa 488), (3) the basal body channel was detected (Alexa 555). The detector gain and the laser intensity for the protein of interest (Alexa 488) channel were constant for all images acquired. On every plate a negative control (no primary rabbit antibody) was included and recorded to confirm that no nonspecific signals were visible in the Alexa 488 channel. Detector gains and laser intensities for the other channels were minimally adjusted between different plates and cell lines, if needed (the different cell lines showed different strength of ARL13B PCNT and DAPI signals and thus, customization was required). Acquisition of images was conducted fully automatically, at least 4 images at random positions per well were acquired. If these images did not contain cilia or were deemed insufficient for deciding on a subcellular location (in the IF image annotation workflow described below), additional images at random positions were acquired. For ASC52telo cells, in this additional acquisition step, the field of views were placed manually in ciliated regions with many cells (without looking at the protein of interest channel).

#### Image processing

Images were exported to OME-TIFF format through Leica LASX’s inbuilt 3D viewer software. The OME-TIFF images were next reprocessed by a custom-written ImageJ plugin (https://github.com/CellProfiling/HPA_Convert_Sp8_To_OMETIF) that complements and corrects metadata in the OME-XML strings in the tiff files based on the xml-based metadata files co-saved from Leicás LAS-X software. Next all images were imported into the Laboratory Internal Management System (LIMS) of the Human Protein Atlas. Here, the intensity range for all channels was adjusted by an automatic script lowering the maximum displayed intensity value to improve visibility of stainings. The maximum display intensity was, however, never allowed to fall below 10,000 a.u., which corresponds to a value where in a negative control staining no signals were visible (Negative control staining was obtained by keeping all markers and secondary antibodies in the procedure but leaving out an antibody for the protein of interest). During manual annotation (see below), the maximum displayed intensity value was - if needed - further adjusted to better visualize ciliary signals but also never allowed to be set below 10,000 a.u. For display on the https://www.proteinatlas.org webpage, all images were converted from TIFF to JPEG file format based on the set maximum display intensity range.

#### Image annotation

The subcellular location of each protein was manually determined based on the signal pattern and relation to the markers for nucleus (DAPI), cilia (ARL13B), and basal bodies (PCNT). 35 different subcellular locations (Intermediate filaments, Actin cytoskeleton, Microtubules, Microtubule ends, Centriolar satellites, Mitochondria, Aggresome, Cytosol, Cytoplasmic bodies, Rods & Rings, ER, Golgi, Vesicles, Peroxisomes, Endosomes, Lysosomes, Lipid droplets, Plasma membrane, Cell Junctions, Focal Adhesions, Nucleoplasm, Nuclear membrane, Nucleoli, Nucleoli Fibrillar center, Nucleoli rim, Nuclear speckles, Nuclear bodies, Micronucleus, Kinetochore, Mitotic chromosome, Cytokinetic bridge, Midbody, Midbody ring, Cleavage furrow, Mitotic spindle), the standard for the subcellular section of the human protein atlas^28^, and four additional cilia-related locations were annotated for all images: primary cilium, basal body, primary cilium tip, primary cilium transition zone. If more than one location was detected, they were defined as the main or additional location depending on the relative signal strength between the location and the most common location when including multiple cell lines, including also non-ciliated cell lines in the HPA. Variation between single cells was annotated as variation in the intensity or spatial distribution based on a visual inspection. The staining was not annotated if considered negative or unspecific.

Of note, given the resolution of the method, it was not possible to distinguish a transition zone and primary cilium staining from an only primary cilium staining unless the transition zone staining is significantly brighter than the whole cilium staining. The proteins annotated to stain the transition zone showed a clear only transition zone staining in at least one cell line. The same applied to ciliary tip proteins.

#### Image analysis

OME-TIFF files for individual channels and slices were fetched from the internal image database of the Human Protein Atlas and composed into z-stack tif-files. To segment individual cell nuclei, the 3D image channel of DAPI staining was blurred using a Gaussian kernel (standard deviation of 3), then we applied a threshold of 0.75 times Otsu threshold to binarize the blurred image. We further performed morphological closing operation with a ball kernel of radius 3 to the binarized image, and filled the internal holes within each foreground region. To segment individual cilia and basal body from the 3-dimensional image stack, we firstly blurred the image using a Gaussian kernel (standard deviation of 1) and subtracted the background which is obtained by blurring the original image using a larger Gaussian kernel (standard deviation of 10). Next, we set threshold values to segment the cilia and basal body objects. We empirically found that Otsu thresholding and 10 times of triangle thresholding values work best for basal body and cilia, respectively. To refine the segmented structures, we performed image opening using a ball kernel of radius 1, then removed small unconnected objects from the segmentation (objects smaller than 75 and 200 pixels for basal body and cilia, respectively). We also removed the falsely segmented cilia that is not attached to any segmented basal bodies.

To analyze how many cells and cilia were in the data set, as well as the ciliation rate, we used a new version of CiliaQ that tracks basal bodies / centrosomes and cilia (CiliaQ Version v0.2.1) ^90^. The number of cells was estimated based on the number of detected basal bodies.

Considering the different sizes of intensity profiles associated with each cilia, we projected each path onto the same 1-dimensional grid from 0 to 1 with 100 intervals (the starting and ending points are mapped to 0 and 1 respectively) and interpolated the profile according to the grid. Specifically, for cilia that are longer than 1.16 micrometers, we partitioned it into two pieces (0 to 1.16 micrometers and 1.16 micrometers to its end) and projected the first part onto the grid of 0 - 20 and the second part onto the grid of 21 - 100. For the cilia shorter than 1.16 micrometers, we projected it onto the grid points 0 to 20 and filled the vector by 0s after that. The rationale here is to prevent stretching protein intensity at the bottom of short cilia too much and to better characterize the non-deformed basal body / transition zone. The stretching creates ciliary intensity profiles representing vectors of a fixed length of 100. All profiles were further normalized by the maximal values among all profiles belonging to the same staining and smoothened each profile using a 1-dimensional Gaussian kernel (standard deviation of 8). To visualize the differences between profiles associated with proteins at different subcellular structures, for example, proteins located at basal body, primary cilium, primary cilium tip and transition zone, we plotted the median, 25 and 75 percentiles among intensity profiles at each grid point along the normalized cilia path for proteins localized in each of those structures.

#### Cell cycle prediction and analysis

To investigate the relationship between nuclear or ciliary protein localization and the cell cycle in RPTEC/TERT1 cells, we formulated the problem as an image-to-image translation task. Specifically, given a set of reference fluorescence channels including nuclear staining, cilia, and basal body markers, we trained a model to predict a corresponding cell cycle marker. Our model is built upon aDPN-UNet architecture ^133^ with three input channels and one output channel.

The model was trained on a dataset of 111 3D images of RPTEC/TERT1 cells stained for nuclei (DAPI), cilia (ARL13B antibody), basal bodies (PCNT), and GMNN (HPA049977, 1:186) generated as described above. 101 images were used for training (containing approximately 2000 cells), and the remaining 10 images were held-out for validation. We converted each 3D multichannel image stack into a 2D image for input into the model as follows: for cell nucleus staining (DAPI) and protein (GMMN) staining, we chose the z-slice with the largest cell nucleus area; for cilia and basal body staining, we performed a maximal intensity projection across all z-slices. Prior to model training, the DAPI, cilia, and basal body images were normalized to values ranging from 0 to 1 based on the 99th percentile of the intensity distribution for all samples in the same well. For the protein (GMNN) channel, intensities were initially clipped at an intensity value of half the bit depth to enhance contrast, and then scaled to 0 to 1.

The model was trained from scratch using a mean squared error (MSE) loss function and an Adam optimizer with a fixed learning rate of 5e-4. Training converged in ∼6 hours, with early stopping triggered after 10 epochs of no improvement in validation loss. After training, the model was used to predict a virtual staining image of the GMNN marker based on the reference inputs. Single-cell predicted nuclear GMNN intensity values were then extracted by determining the average intensity within the nucleus using the nuclear masks.

To further validate the model, the prediction was also applied to the held-out images (10 3D images featuring 190 cells) and stainings for the other cell cycle markers CCNA2 (Antibody: HPA020626, 1:162; five 3D images yielding 136 cells), CDT1 (HPA017224, 1:9; 26 3D images yielding 508 cells), or KIF18B (HPA024205, 1:43; five 3D images yielding 103 cells). From the original image stacks and the predicted GMNN channel, predicted nuclear GMNN intensity as well as ground-truth nuclear intensity of the stained protein was determined. Predicted and ground truth intensity values were each percentile normalized on the staining level (1% percentile represents the value 0, 99% percentile represents the value 1) and then plotted (Figure 4H).

To determine the relationship of predicted nuclear GMNN intensity and ciliary CKAP2 intensity at single cell level, the prediction model was applied to the 3D images featuring CKAP2 staining processed into 2D images as for training (Atlas data set: four 3D images featuring 91 cells; Further validation set: 150 3D images featuring 4,149 cells) and predicted nuclear GMNN intensity and the center x,y coordinates of each nucleus were determined. Predicted nuclear GMNN intensity values were percentile-normalized on the staining level (1% percentile represents the value 0, 99% percentile represents the value 1). The same images underwent CiliaQ analysis to determine ciliary parameters like basal body x and y center and ciliary centerline average intensity in the CKAP2 channel. Next, each nucleus was matched to a basal body by computing a distance matrix and applying linear sum assignment for optimal (minimum distance) pairing. The Euclidean distance matrix, before linear sum assignment, was modified as follows: 1.) All distances were clipped to 250 px (ca. 17.5 µm) to avoid that missing pairs (no nucleus or no basal body detected) distorted the assignment through introducing unreasonable long-range assignments that largely influence the sum of assigned distances that is optimized in the linear sum assignment. 2.) The distance matrix was squared to enforce preference for short range over long range assignments. After performing linear sum assignment, all nuclei with basal body to nucleus distance above 250 px (17.5 µm) were removed to exclude random assignments made by the algorithm for out-of-range partners. For plotting (Figures 4I and 4J) only cells with cilia were considered and the ciliary CKAP2 intensity was percentile normalized (0 = 1%, 1 = 99%).

#### Human tissue sample preparation

The tissue sample preparation protocol used was the same as that described earlier by Danielsson et al. ^134^. Tissue MicroArrays (TMA) were constructed with strategies used in the Human Protein Atlas previously described ^135,136^, in Formalin-fixed paraffin-embedded (FFPE) format. In summary, hematoxylin-eosin stained tissue sections from FFPE donor blocks were examined to confirm histology and identify representative regions for sampling into the TMA. Normal tissue was characterized as microscopically normal, predominantly sourced from proximity areas to surgically excised tumors. TMA blocks were created, each containing triplicate 1 mm cores from the different types of normal tissue.

#### Immunofluorescence staining of tissues

We used the same indirect immunofluorescence staining protocol described by Martinez Casals et al. ^137^. with some adjustments. In brief: single section from each of the human TMA FFPE blocks was sectioned at 4 µm thickness. Each section was dewaxed and underwent heat-induced epitope retrieval (HIER) treatment using pH 9 EDTA buffer (Akoya Biosciences, AR900250ML) in a pressure cooker (Bio SB TintoRetriever, BSB-7087) at 114–121 °C during 20 minutes. After two ddH2O washes, the material was exposed to a photobleaching procedure to reduce tissue autofluorescence; following a blocking step using TNB buffer containing 0.1 M Tris-HCl /0.15 M NaCl/0.5% blocking reagent pH 7.5 (Akoya, SKU FP1020) for 30 min. Primary antibodies targeting ARL13B (Abcam, ab136648, RRID:AB_3073658, mouse IgG2a, 1:300), PCNT (Abcam, ab28144, RRID:AB_2160664, mouse IgG1, 1:500), and CFAP300 (HPA, HPA038585, rabbit polyclonal, 1:100) were diluted in antibody diluent containing 0.3 % Triton (Sigma Aldrich, T8787) 1x PBS pH 7.4 and added on the TMA section to be incubated overnight at 4°C. The next day, the material was washed with TBS-T and blocked using TNB for 30 min. Secondary antibodies, goat-anti-mouse-IgG2a-Cross-Adsorbed-Alexa647 (ThermoFisher, A-21241, 1:600), goat-anti-mouse-IgG1-Cross-Adsorbed-Alexa555 (ThermoFisher, A-21127, 1:600), goat-anti-rabbit-IgG (H+L)-Highly-Cross-Adsorbed-Alexa488 (Invitrogen, A11034, 1:600) were diluted in TNB, containing Hoechst (ThermoFisher, H3570), and added on the section for 90 min incubation at room temperature. Lastly, the stained material was mounted with Fluoromount-G (Thermo Fisher Scientific, 00-4958-02) to proceed with image acquisition.

#### Confocal microscopy of tissue microarrays

Regions of interest in the TMA were image acquired using Leica STELLARIS 8 confocal microscope equipped with a 63 HC PL APO 63x/1.20 water-immersion CS2 objective (Leica Microsystems. Mannheim, Germany). The settings for each image were Pinhole 1 Airy unit, 16-bit acquisition, 1x zoom, 2048x2048 pixels field of view, Z-stack size 9.1 µm, number of steps 14, resulting in a pixel size of 0.09 μm and a z-step size of 0.7 μm. Detectors used: HyD S1 was used for Hoechst channel, HyD H2 for A488 channel, Hyd S3 regarding the A555 channel, and Hyd X4 for Alexa 647 channel. The “Lightning” deconvolution software (Leica Microsystems, Mannheim, Germany), which was integrated into the microscope software, was used to deconvolve the images after recording.

#### Immunohistochemistry images

The immunohistochemical stainings were performed as part of the regular Human Protein Atlas workflow, as described elsewhere. ^27,35^

#### Exploration of clinical databases

Clinical samples analyzed at the Karolinska University Hospital by whole genome sequencing (WGS) were searched for interesting variants in genes who were classified in OMIM ^102^ as without a phenotype or not annotated in OMIM at all (Figure 6A, Table S7). The overall detailed analytical process of WGS has previously been described in Stranneheim et al. ^120^ and in Ek et al. ^121^ Clinical WGS data from three groups of patients were queried: 1) patients with a suspected ciliopathy disorder without a causative variant found in a clinical WGS (n=157), 2) fetuses with an ultrasound-detected malformation without a causative variant found in a clinical WGS (n=470) and 3) patients who had undergone a clinical patient-parental trio WGS analysis without a causative variant found (n>1,500). We searched for potentially clinically relevant single nucleotide variants in the WGS data from these three patient groups by one gene at a time, filtering for the found variants so that only frameshift variants, nonsense variants and predicted splicing variants in a homozygous, compound heterozygous or *de novo* state were retained.

#### Proximity ligation assay

Cells were seeded and fixed as described above under Immunocytochemistry. Cells were washed with TBS and permeabilized with 0.5% Triton X-100 (Sigma Aldrich) in TBS for 3 × 5 min. Next, the NaveniFlex Cell Atto647N kit (Navinci Diagnostics, NC.MR.100 Atto647N) was applied according to the manufacturer’s instructions. As a bait, a murine ARL13B antibody was used (Abcam, ab136648, mouse IgG2a, 1:2000). As prey, the HPA antibodies for CREB3 (Human Protein Atlas, HPA077421, 1:64) or GPR75 (Human Protein Atlas, HPA056032, 1:143) were used. As positive control, a rabbit-derived ARL13B antibody was used (Proteintech, 17711-1-AP, 1:1000). As negative controls, no rabbit primary antibody and a rabbit antibody targeting the nuclear protein PDS5A (Human Protein Atlas, HPA036661, 1:86) were used. After application of the NaveniFlex kit, wells were washed 1x with TBS and post-stained to label the individual antibodies as follows: Secondary antibodies, goat-anti-rabbit-AlexaFluorPlus-488 (1:800, Invitrogen, A32731) and goat-anti-mouse-Alexa-555 (1:800, Invitrogen, A21424), diluted in blocking buffer (TBS + 4% FBS) were added and the plates were incubated for 90 min at room temperature. Next, cells were washed 4 x 5 min with TBS. Before and after post-staining, cells were briefly investigated with a high-resolution confocal microscope (Leica STELLARIS 8) to validate that post-staining did not alter visibility of proximity ligation signals. Finally, images were recorded with the Leica STELLARIS 8 microscope, applied as described in the section “Confocal microscopy of tissue microarrays” but without applying the “Lightning” deconvolution software.

#### Heterologous expression of tagged proteins

CaMK1G-HaloTag and alphaTag-CREB3 vectors on the all-in-one Tet3G backbone ^138^ (XLone-GFP, gift from Xiaojun Lian, Addgene plasmid # 96930; http://n2t.net/addgene:96930; RRID:Addgene_96930)) were synthesized by VectorBuilder. hTERT-RPE cells (RPE-1, ATCC CRL-4000) were electroporated with the hyperactive piggyBac transposase vector (VectorBuilder, Inc) plus either the CaMK1G-HaloTag or alphaTag-CREB3 vector using Lonza nucleofector in P3 solution (Lonza Bioscience) and grown in DMEM:F12 media (ATCC-30-2006) with 10% Tet-free FBS (Gemini). The cells were then selected by blasticidin to create stable cells.

Tet-on CAMK1G-HaloTag and alphaTag-CREB3 RPE-1 cells were plated in 35 mm coverslip bottom dishes (Cellvis) in 10% FBS DMEM:F12 media (ThermoFisher Scientific). The next day, the cells were serum deprived with 0% FBS DMEM:F12 medium with 500 ng/ml doxycycline to induce transgene expression. After 48 hours of starvation, CAMK1G-HaloTag cells were first stained with Janelia Fluor 552 (JF-552)HTL (250 nM, 30 min) and then both theCAMK1G-HaloTag and alphaTag-CREB3 cells were fixed with 4% paraformaldehyde (PFA) for 15 minutes at room temperature, followed by three washes with phosphate-buffered saline (PBS). Permeabilization was performed using 0.02% Triton X-100 in PBS for 15 minutes. Cells were then blocked in BlockAid (Thermo Fisher Scientific, B10710) for 15 minutes at room temperature. Primary antibody incubation was performed overnight at 4°C using Rat anti-ARL13b (BiCell, Cat# 90413h) diluted 1:100 in BlockAid. The following day, cells were washed three times with PBS and incubated for 1 hour at room temperature with secondary antibodies diluted in BlockAid: anti-AlphaTag-Alexa647 (for alphaTag-CREB3 cells only, 1:100, NanoTag Biotechnologies, Cat# N1502-AF647-L) and Alexa Fluor 488-conjugated anti-Rat IgG (Invitrogen, Cat# A48269, 1:1000). After three final washes with PBS, samples were prepared for imaging.

Images were acquired on a Leica Stellaris 8 confocal microscope using a 40x glycerol immersion objective (NA 1.25, HC PL APO CS2). Z-stacks were collected in xyz mode with a z-galvo scanner, capturing 22 optical sections at 0.36 µm intervals. Images were recorded at 1832 × 1832 resolution with a pixel size of 0.08 µm, using a 67.9 µm pinhole (equivalent to 1 Airy unit). For CaMK1G-HaloTag cells, the eExcitation was performed with 488 nm (1% power) and 561 nm (7% power) lasers, and fluorescence was detected using HyD detectors with emission ranges of 499– 556 nm for ATTO 488 and 570-650 nm for JF552. For alpha-CREB3 cells, .excitation was performed with 488 nm (1% power) and 638 nm (7% power) lasers, and fluorescence was detected using HyD detectors with emission ranges of 499–556 nm for ATTO 488 and 657–793 nm for Alexa-647. Acquisition was done in photon counting mode with a pixel dwell time of 0.775 µs and two-line accumulation. Images were processed using LAS X and FIJI/ImageJ.

### Quantification and statistical analysis

#### Hierarchical clustering of intensity profiles

Single-cilia intensity profiles were aggregated to mean ciliary intensity profiles per protein and cell line utilizing Python v3.10. Each average profile was scaled by dividing all intensity values by the maximum intensity value of that profile. Hierarchical clustering of the aggregated profiles was conducted for each cell line, employing the pdist() and linkage() functions from the SciPy library (v1.14.0) ^139^ with Euclidean distance and the Unweighted Pair Group Method with Arithmetic Mean (UPGMA). Clusters were identified using the AgglomerativeClustering() function from scikit-learn v1.5.1 ^140^, with a distance threshold chosen to reflect the similarity matrix of the clustered profiles. Finally, the clustered average intensity profiles were visualized using the clustermap() and heatmap() functions from the seaborn library v0.13.2. Additionally, a functional enrichment analysis for each cluster was performed using the G:Profiler Python library v1.0.0 ^141^. GO Biological Processes and GO Molecular Functions ontologies were used, the human genome was set as background, g:SCS algorithm was used to compute multiple testing corrections for p-values and a significance threshold of p<0.05 was selected. The resulting significant terms were simplified by removing all terms that were significant for more than one cluster per cell line.

#### Multilocalization analysis

To assess whether ciliary proteins localize to significantly more subcellular compartments than the overall proteome, we first counted the number of subcellular compartments annotated for each protein in the Human Protein Atlas. We then compared the distribution of the number of subcellular localization annotations for all proteins in the HPA ( with any subcellular location annotation) to that of ciliary proteins—specifically, those annotated with "primary cilium tip," "primary cilium," and "primary cilium transition zone." A two-sided Mann-Whitney U test was used to determine statistical significance, with a P-value threshold set at P < 0.01. Version v24 of the Human Protein Atlas was used.

To study the locations, where multilocalizing proteins were observed, the annotations on the staining level (data split by protein and cell line) from all cell lines were converted into edges with the python package networkx (version 3.2) and next plotted using the python package d3blocks (version 1.4.6).

#### Functional enrichment analyses

Functional enrichment analyses were conducted in R v4.4.1 using clusterProfiler v4.12.6 ^142^, enrichplot v1.24.4, and simplifyEnrichment v1.14.1 ^143^. Overrepresentation of Gene Ontology Biological Process terms compared to all human genes was computed using the compareCluster function of clusterProfiler with org.Hs.eg.db as annotation database. The results were controlled for multiple hypothesis testing by calculating adjusted P values using the Benjamini-Hochberg Procedure and q values and restricting the results to adjusted P values < 0.01 and q values < 0.01. Results were visualized using the dotplot function from enrichplot. Additionally, the resulting list of significantly enriched GO terms for each variable in each comparison was simplified by calculating and clustering the corresponding semantic similarity matrix of the significant GO terms. Relational similarity was used as the similarity measure and binary_cut as clustering method. For each of the resulting clusters, a word cloud was drawn summarizing the biological functions with keywords in every GO term cluster. For this, the enrichment of keywords compared to the background GO vocabulary is determined using Fisher’s Exact test and mapped to the fontsize of the keywords in the word clouds. Overrepresentation of Reactome pathways was determined equivalently, but were not further simplified.

To compare ciliary locations to cellular locations the 35 HPA locations were simplified into four groups: cytoplasm, membrane, nucleus and mitotic. Cytoplasm collected annotations for Intermediate filaments, Actin cytoskeleton, Microtubules, Microtubule ends, Centriolar satellites, Mitochondria, Aggresome, Cytosol, Cytoplasmic bodies, Rods & Rings. Membrane contained the endo-membrane system, collecting annotations for ER, Golgi, Vesicles, Peroxisomes, Endosomes, Lysosomes, Lipid droplets, Plasma membrane, Cell Junctions, Focal Adhesions. Nucleus collected annotations for Nucleoplasm, Nuclear membrane, Nucleoli, Nucleoli Fibrillar center, Nucleoli rim, Nuclear speckles, Nuclear bodies, Micronucleus. Mitotic collected annotations for Kinetochore, Mitotic chromosome, Cytokinetic bridge, Midbody, Midbody ring, Cleavage furrow, Mitotic spindle.

### Additional resources

Human Protein Atlas: https://www.proteinatlas.org/

## Supplemental information

Document S1: Figures S1–S9 and Note S1.

Table S1. Excel file listing the genes appearing in existing datasets and databases (see Star Methods), which of these genes were among the 1,992 candidates whose proteins we stained, and the list of 715 genes whose proteins we detected at cilia, in which of the four compartments we detected the proteins, and the overlap of these genes to genes listed in existing datasets and databases, too large to fit in a PDF, related to Figures 1I, 2F, and 2G.

Table S2. Excel file containing functional enrichment results of the comparison between primary cilium + tip and basal body + transition zone, including clusters of the GO term similarity clustering, too large to fit in a PDF, related to Figure 3A and S1.

Table S3. Excel file containing functional enrichment results of the comparison between cell lines including clusters of GO term similarity clustering, too large to fit in a PDF, related to Figure 4C and S2.

Table S4. Excel file containing functional enrichment results of the comparison between proteins with stable intensity and proteins with variable intensity between individual cells including clusters of GO term similarity clustering, too large to fit in a PDF, related to Figure 4D and S3.

Table S5. Excel file containing the comparative GO BP enrichment results comparing the four ciliary locations (cilia tip, primary cilia excluding tip, transition zone and basal body excluding transition zone) as well as the rest of the cell (cytoplasm, membrane, nucleus and mitotic), too large to fit in a PDF, related to Figures S4.

Table S6. Excel file showing the proteins along with their functional enrichment analysis results for each cluster derived from hierarchically clustering ciliary intensity profiles, averaged by protein and cell line (as shown in Figure 5), too large to fit in a PDF, related to Figure 5.

Table S7. Excel file classifying the genes into three groups based on OMIM annotations (see Star Methods), too large to fit in a PDF, related to Figure 6A.

Table S8. Excel file listing the HPA antibodies applied in this study for the 715 genes whose proteins were detected in cilia. Validation scores for Immunocytochemistry (ICC), Immunohistochemistry (IHC), Western Blot (WB), and Protein Array (PA) based validation are provided based on HPA v24. Too large to fit in a PDF, related to Figure 2.

