## Supplementary material for "Intrinsic Heterogeneity of Primary Cilia Revealed Through Spatial Proteomics": Document S1

### SUPPLEMENTARY INFORMATION
### (Document S1)

### FOR

### Intrinsic Heterogeneity of Primary Cilia Revealed Through Spatial Proteomics

Jan N. Hansen [1,2], Huangqingbo Sun [2,†], Konstantin Kahnert [2,†], Eini Westenius [3,4], Alexandra Johannesson [1], Carmela Villegas [5], Trang Le [1,2], Kalliopi Tzavlaki [1], Casper Winsnes [1], Emmie Pohjanen [1], Anna Mäkiniemi [1], Jenny Fall [1], Frederic Ballllosera Navarro [2], Anna Bäckström [1,2], Cecilia Lindskog [6], Fredric Johansson [1], Kalle von Feilitzen [1], Angelica M. Delgado-Vega [3,4], Anna Martinez Casals [2], Diana Mahdessian [1], Mathias Uhlén [1,7], Shu-Hsien Sheu [5], Anna Lindstrand [3,4], Ulrika Axelsson [1], Emma Lundberg [1,2,8,9,*]

1 Science for Life Laboratory, School of Engineering Sciences in Chemistry, Biotechnology and Health, KTH - Royal Institute of Technology, Stockholm, Sweden.
2 Department of Bioengineering, Stanford University, Stanford, CA, USA.
3 Department of Molecular Medicine and Surgery, Karolinska Institutet, Stockholm, Sweden.
4 Department of Clinical Genetics and Genomics, Karolinska University Hospital, Stockholm, Sweden.
5 Chan Zuckerberg Imaging Institute, Redwood City, CA, USA
6 Department of Immunology, Genetics and Pathology, Cancer Precision Medicine Research Unit, Uppsala University, Uppsala, Sweden.
7 Department of Neuroscience, Karolinska Institutet, Stockholm, Sweden.
8 Department of Pathology, Stanford University, Stanford, CA, USA.
9 Chan Zuckerberg Biohub, San Francisco, San Francisco, CA, USA.
[†] Contributed equally

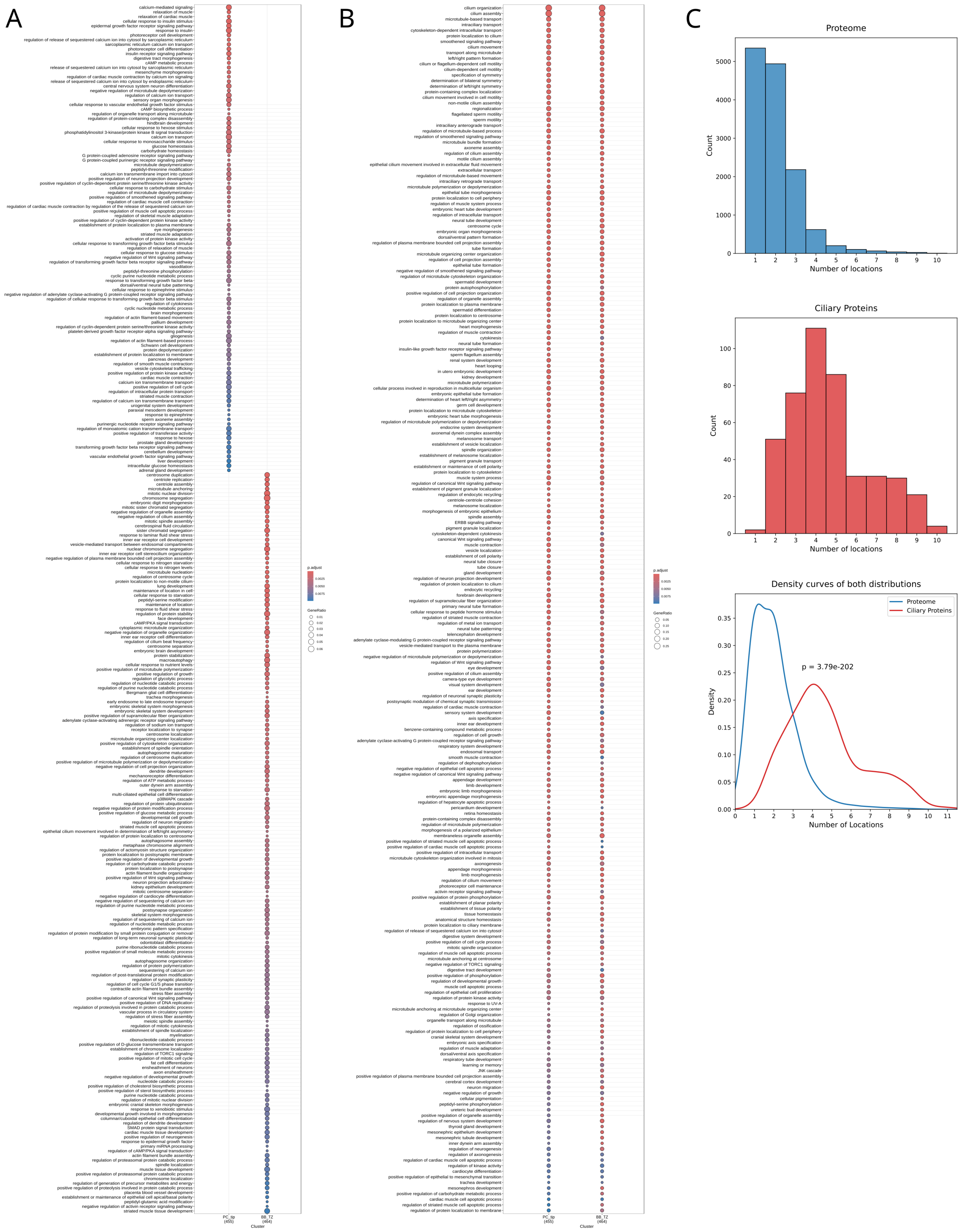
A
B
C
Proteome
Count
Number of locations
Ciliary Proteins
Count
Number of locations
Density curves of both distributions
Proteome
Ciliary Proteins
p = 3.79e-202
Density
Number of Locations
p.adjust
GeneRatio
Cluster

#### Figure S1:

Dotplots showing all significantly enriched GO BP terms in primary cilia including tip, and basal body including transition zone. **A)** shows terms only significantly enriched for one of the locations and **B)** shows terms significantly enriched in both locations. Dot size shows the ratio of identified proteins of a given term compared to all proteins of the term whereas the dot color shows the Benjamini-Hochberg adjusted P-value of the overrepresentation analysis. **C)** Top, distribution of the number of subcellular location annotations of all proteins in the HPA. Middle, distribution of the number of subcellular location annotations of all ciliary proteins. Bottom, density curves of the two distributions along with Mann-Whitney U test derived P-value showing that ciliary proteins localize to significantly more subcellular compartments compared to the proteome in general.

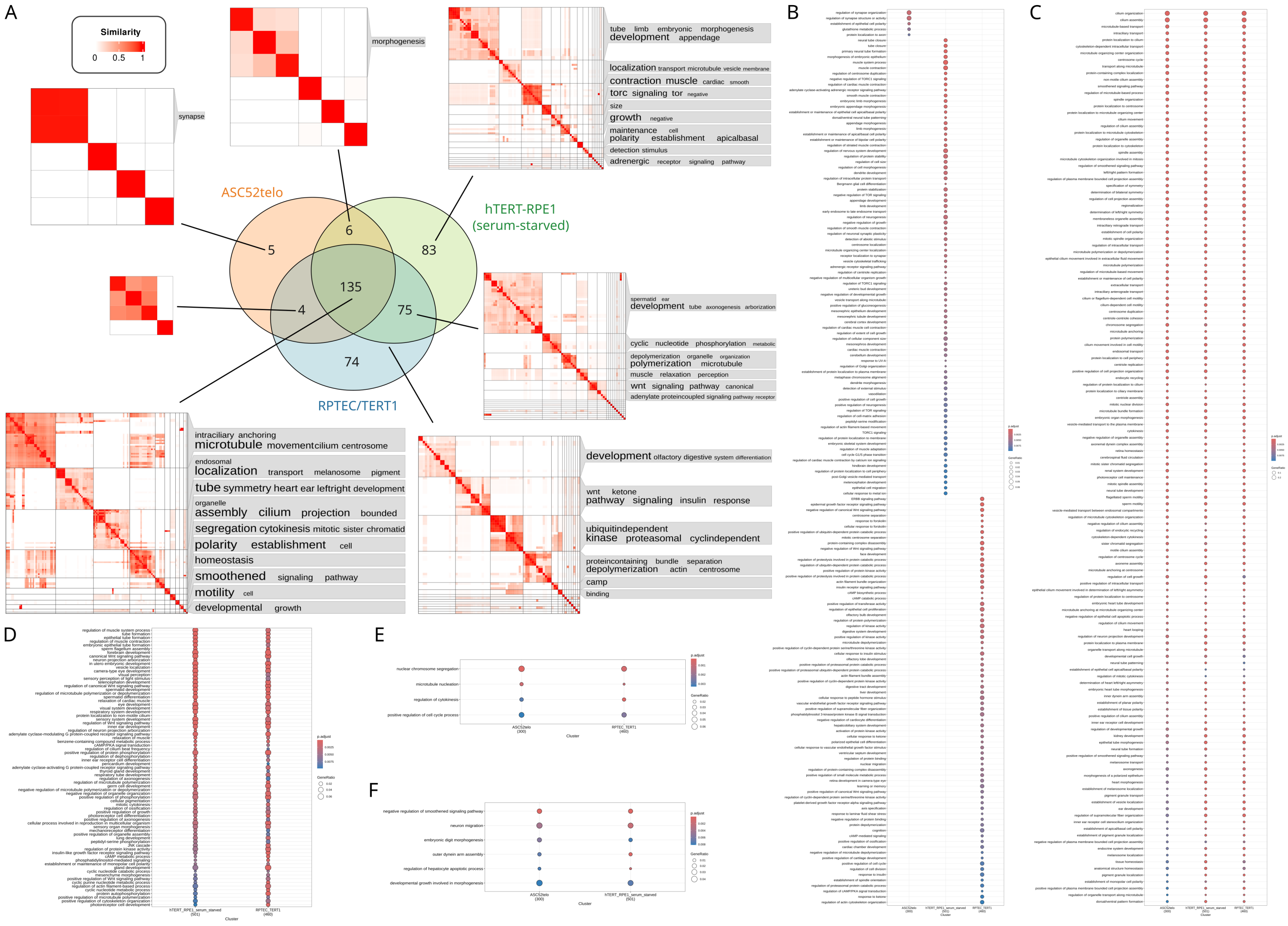


### Figure S2:

**A)** Comparative Gene Ontology enrichment analysis of Biological Process terms between the three included cell lines to investigate cell-line differences and commonalities. The Venn diagram shows the overlap of significantly enriched terms (adj. P value < 0.01 and q value < 0.01) for each cell line. Significant terms of each section of the Venn diagram were clustered by semantic similarity, and for each resulting GO term cluster, a word cloud is shown summarizing the biological functions of the terms in the cluster based on keywords enrichment of term names. The font size in the word clouds correlates with -log(P value) of the key word enrichment, i.e. the larger, the more enriched a keyword is compared against background GO vocabulary. The color in the heat maps represents the semantic similarity. **B-F)** Dotplots showing all significantly enriched GO BP terms in ASC52telo, hTERT-RPE1 and RPTEC/TERT1 cells. B**)** shows terms only significantly enriched for one of the cell lines, C) shows terms significantly enriched in all three cell lines, D**)** terms significantly enriched in both hTERT-RPE1 and RPTEC/TERT1 cells but not ASC52telo, E**)**

terms significantly enriched in both ASC52telo and RPTEC/TERT1 cells but not hTERT-RPE1, F**)** terms significantly enriched in both ASC52telo and hTERT-RPE1 cells but not RPTEC/TERT1. Dot size shows the ratio of identified proteins of a given term compared to all proteins of the term whereas the dot color shows the Benjamini-Hochberg adjusted P-value of the overrepresentation analysis.

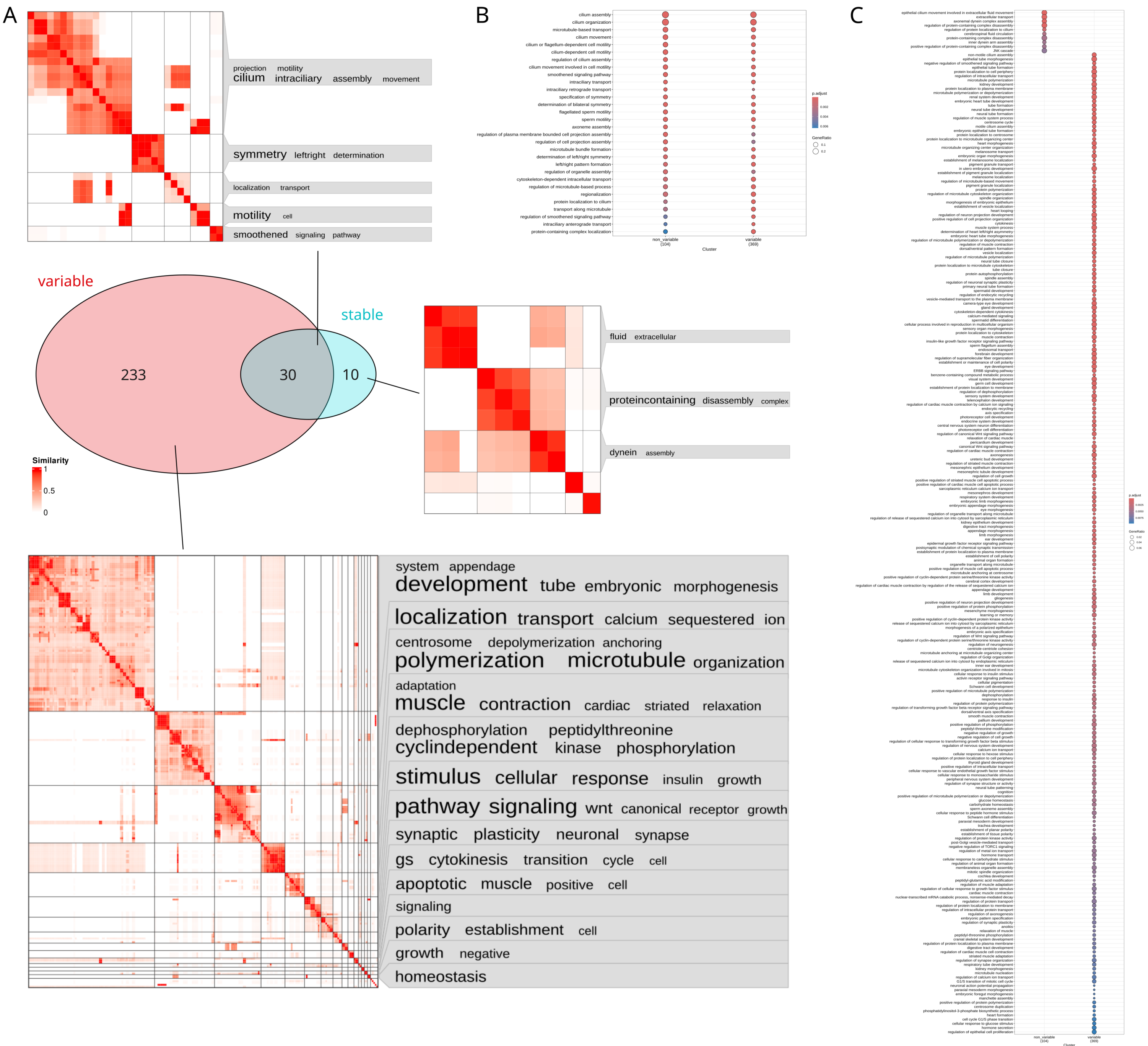


### Figure S3:

**A)** Comparative Gene Ontology enrichment analysis of GO BP terms of proteins annotated as displaying intensity variation in one of the four ciliary locations or annotated as stable. The Venn diagram shows the number of terms found to be significantly enriched (adj. P value < 0.01 and q value < 0.01) for variable proteins (n=199) or non-variable proteins (n=15), as well as for both (n=33). Significant terms were clustered by semantic similarity, and for each resulting GO term cluster, a word cloud is shown summarizing the biological functions of the terms in the cluster based on keywords enrichment of term names. The font size in the word clouds correlates with

-log(P value) of the key word enrichment, i.e. the larger, the more enriched a keyword is compared against background GO vocabulary. The color in the heat maps represents the semantic similarity. **B-C)** Dotplots showing all significantly enriched GO BP terms of proteins annotated as displaying intensity variation in one of the four ciliary locations or annotated as stable. B) shows terms significantly enriched for both variable and stable proteins and C) shows terms only significantly enriched for either variable or stable proteins. Dot size shows the ratio of identified proteins of a given term compared to all proteins of the term whereas the dot color shows the Benjamini-Hochberg adjusted P-value of the overrepresentation analysis.

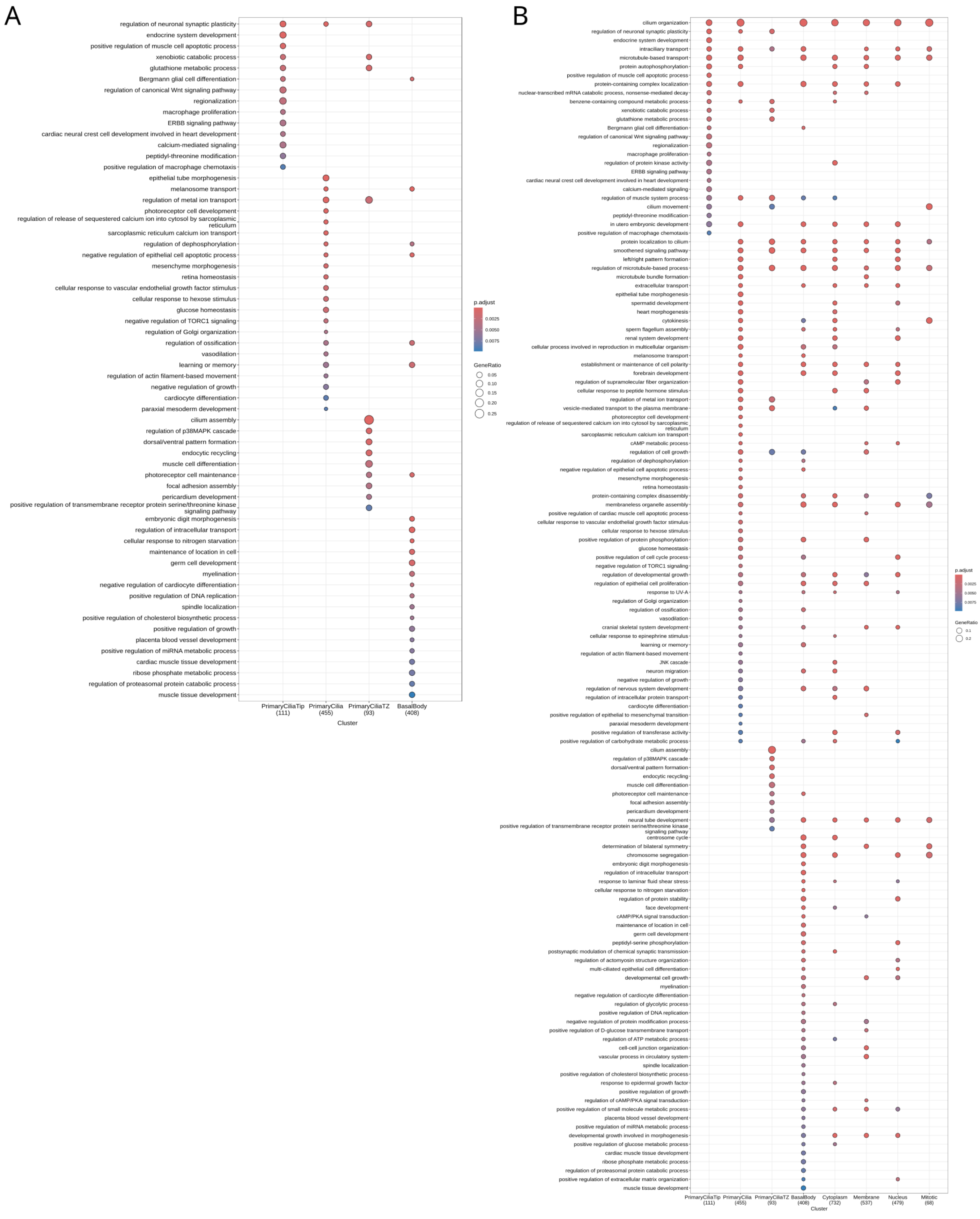
A
regulation of neuronal synaptic plasticity
endocrine system development
positive regulation of muscle cell apoptotic process
xenobiotic catabolic process
glutathione metabolic process
Bergmann glial cell differentiation
regulation of canonical Wnt signaling pathway
regionalization
macrophage proliferation
ERBB signaling pathway
cardiac neural crest cell development involved in heart development
calcium-mediated signaling
peptidyl-threonine modification
positive regulation of macrophage chemotaxis
epithelial tube morphogenesis
melanosome transport
regulation of metal ion transport
photoreceptor cell development
regulation of release of sequestered calcium ion into cytosol by sarcoplasmic reticulum
sarcoplasmic reticulum calcium ion transport
regulation of dephosphorylation
negative regulation of epithelial cell apoptotic process
mesenchyme morphogenesis
retina homeostasis
cellular response to vascular endothelial growth factor stimulus
cellular response to hexose stimulus
glucose homeostasis
negative regulation of TORC1 signaling
regulation of Golgi organization
regulation of ossification
vasodilation
learning or memory
regulation of actin filament-based movement
negative regulation of growth
cardiocyte differentiation
paraxial mesoderm development
cilium assembly
regulation of p38MAPK cascade
dorsal/ventral pattern formation
endocytic recycling
muscle cell differentiation
photoreceptor cell maintenance
focal adhesion assembly
pericardium development
positive regulation of transmembrane receptor protein serine/threonine kinase signaling pathway
embryonic digit morphogenesis
regulation of intracellular transport
cellular response to nitrogen starvation
maintenance of location in cell
germ cell development
myelination
negative regulation of cardiocyte differentiation
positive regulation of DNA replication
spindle localization
positive regulation of cholesterol biosynthetic process
positive regulation of growth
placenta blood vessel development
positive regulation of miRNA metabolic process
cardiac muscle tissue development
ribose phosphate metabolic process
regulation of proteasomal protein catabolic process
muscle tissue development
PrimaryCiliaTip (111)
PrimaryCilia (455)
PrimaryCiliaTZ (93)
BasalBody (408)
Cluster
p.adjust
0.0025
0.0050
0.0075
GeneRatio
0.05
0.10
0.15
0.20
0.25
B
cilium organization
regulation of neuronal synaptic plasticity
endocrine system development
intraciliary transport
microtubule-based transport
protein autophosphorylation
positive regulation of muscle cell apoptotic process
protein-containing complex localization
nuclear-transcribed mRNA catabolic process, nonsense-mediated decay
benzene-containing compound metabolic process
xenobiotic catabolic process
glutathione metabolic process
Bergmann glial cell differentiation
regulation of canonical Wnt signaling pathway
regionalization
macrophage proliferation
regulation of protein kinase activity
ERBB signaling pathway
cardiac neural crest cell development involved in heart development
calcium-mediated signaling
regulation of muscle system process
cilium movement
peptidyl-threonine modification
in utero embryonic development
positive regulation of macrophage chemotaxis
protein localization to cilium
smoothened signaling pathway
left/right pattern formation
regulation of microtubule-based process
microtubule bundle formation
extracellular transport
epithelial tube morphogenesis
spermatid development
heart morphogenesis
cytokinesis
sperm flagellum assembly
renal system development
cellular process involved in reproduction in multicellular organism
melanosome transport
establishment or maintenance of cell polarity
forebrain development
regulation of supramolecular fiber organization
cellular response to peptide hormone stimulus
regulation of metal ion transport
vesicle-mediated transport to the plasma membrane
photoreceptor cell development
regulation of release of sequestered calcium ion into cytosol by sarcoplasmic reticulum
sarcoplasmic reticulum calcium ion transport
cAMP metabolic process
regulation of cell growth
regulation of dephosphorylation
negative regulation of epithelial cell apoptotic process
mesenchyme morphogenesis
retina homeostasis
protein-containing complex disassembly
membraneless organelle assembly
positive regulation of cardiac muscle cell apoptotic process
cellular response to vascular endothelial growth factor stimulus
cellular response to hexose stimulus
positive regulation of protein phosphorylation
glucose homeostasis
positive regulation of cell cycle process
negative regulation of TORC1 signaling
regulation of developmental growth
regulation of epithelial cell proliferation
response to UV-A
regulation of Golgi organization
regulation of ossification
vasodilation
cranial skeletal system development
cellular response to epinephrine stimulus
learning or memory
regulation of actin filament-based movement
JNK cascade
neuron migration
negative regulation of growth
regulation of nervous system development
regulation of intracellular protein transport
cardiocyte differentiation
positive regulation of epithelial to mesenchymal transition
paraxial mesoderm development
positive regulation of transferase activity
positive regulation of carbohydrate metabolic process
cilium assembly
regulation of p38MAPK cascade
dorsal/ventral pattern formation
endocytic recycling
muscle cell differentiation
photoreceptor cell maintenance
focal adhesion assembly
pericardium development
neural tube development
positive regulation of transmembrane receptor protein serine/threonine kinase signaling pathway
centrosome cycle
determination of bilateral symmetry
chromosome segregation
embryonic digit morphogenesis
regulation of intracellular transport
response to laminar fluid shear stress
cellular response to nitrogen starvation
regulation of protein stability
face development
cAMP/PKA signal transduction
maintenance of location in cell
germ cell development
peptidyl-serine phosphorylation
postsynaptic modulation of chemical synaptic transmission
regulation of actomyosin structure organization
multi-ciliated epithelial cell differentiation
developmental cell growth
myelination
negative regulation of cardiocyte differentiation
regulation of glycolytic process
positive regulation of DNA replication
negative regulation of protein modification process
positive regulation of D-glucose transmembrane transport
regulation of ATP metabolic process
cell-cell junction organization
vascular process in circulatory system
spindle localization
positive regulation of cholesterol biosynthetic process
response to epidermal growth factor
positive regulation of growth
regulation of cAMP/PKA signal transduction
positive regulation of small molecule metabolic process
placenta blood vessel development
positive regulation of miRNA metabolic process
developmental growth involved in morphogenesis
positive regulation of glucose metabolic process
cardiac muscle tissue development
ribose phosphate metabolic process
regulation of proteasomal protein catabolic process
positive regulation of extracellular matrix organization
muscle tissue development
PrimaryCiliaTip (111)
PrimaryCilia (455)
PrimaryCiliaTZ (93)
BasalBody (408)
Cytoplasm (732)
Membrane (537)
Nucleus (479)
Mitotic (68)
Cluster
p.adjust
0.0025
0.0050
0.0075
GeneRatio
0.1
0.2

#### Figure S4:

**A-B)** Dotplot showing a subset of the simplified comparative GO BP enrichment results comparing the four ciliary locations (cilia tip, primary cilia excluding tip, transition zone and basal body excluding transition zone) as well as the rest of the cell (35 HPA annotations grouped into cytoplasm, membrane, nucleus and mitotic, see materials and methods for details). The full list of enriched terms, as well as the simplified list of terms can be found in Table S5. Enrichment results were simplified by clustering the GO terms by semantic similarity (cutoff=0.6) and selecting the most significant term as representative term for the cluster. Dot size shows the ratio of identified proteins of a given term compared to all proteins of the term whereas the dot color shows the Benjamini-Hochberg adjusted P-value of the overrepresentation analysis. A) This plot shows only terms that are only enriched in any of the four cilium locations but not any of the locations in the rest of the cell. **B)** This plot shows only terms that are enriched for at least one ciliary location.

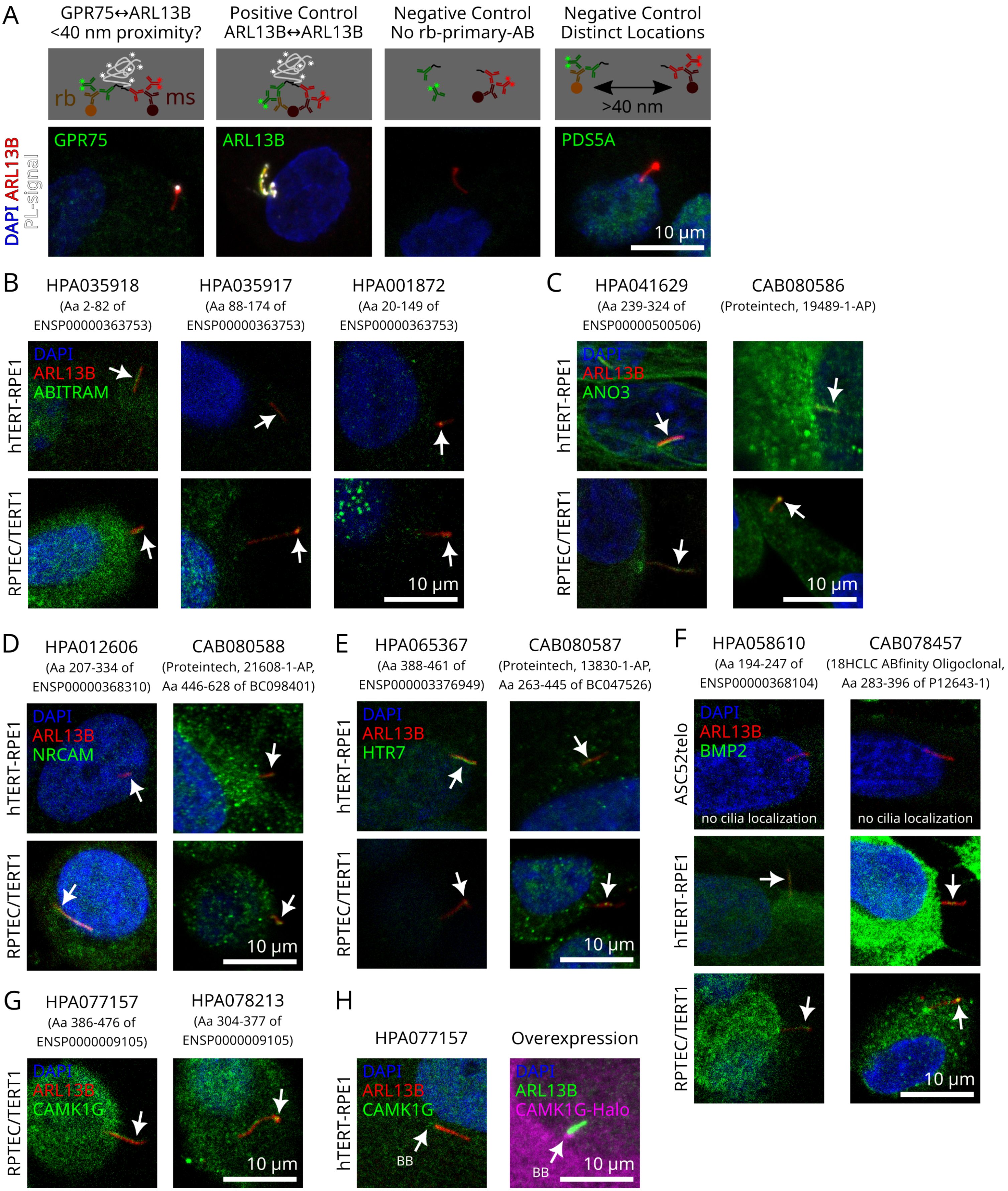
A
GPR75↔ARL13B
<40 nm proximity?
Positive Control
ARL13B↔ARL13B
Negative Control
No rb-primary-AB
Negative Control
Distinct Locations
rb
ms
>40 nm
DAPI ARL13B PL-signal
GPR75
ARL13B
PDS5A
10 µm
B
HPA035918
(Aa 2-82 of ENSP00000363753)
HPA035917
(Aa 88-174 of ENSP00000363753)
HPA001872
(Aa 20-149 of ENSP00000363753)
hTERT-RPE1
RPTEC/TERT1
DAPI
ARL13B
ABITRAM
10 µm
C
HPA041629
(Aa 239-324 of ENSP00000500506)
CAB080586
(Proteintech, 19489-1-AP)
DAPI
ARL13B
ANO3
10 µm
D
HPA012606
(Aa 207-334 of ENSP00000368310)
CAB080588
(Proteintech, 21608-1-AP, Aa 446-628 of BC098401)
DAPI
ARL13B
NRCAM
10 µm
E
HPA065367
(Aa 388-461 of ENSP000003376949)
CAB080587
(Proteintech, 13830-1-AP, Aa 263-445 of BC047526)
DAPI
ARL13B
HTR7
10 µm
F
HPA058610
(Aa 194-247 of ENSP00000368104)
CAB078457
(18HCLC ABfinity Oligoclonal, Aa 283-396 of P12643-1)
ASC52telo
DAPI
ARL13B
BMP2
no cilia localization
no cilia localization
10 µm
G
HPA077157
(Aa 386-476 of ENSP0000009105)
HPA078213
(Aa 304-377 of ENSP0000009105)
DAPI
ARL13B
CAMK1G
10 µm
H
HPA077157
Overexpression
DAPI
ARL13B
CAMK1G
BB
DAPI
ARL13B
CAMK1G-Halo
BB
10 µm

#### Figure S5:

**A)** Example images from a proximity ligation assay revealing that GPR75 and ARL13B localize within 40 nm proximity at cilia of RPTEC/TERT1 cells (see STAR Methods). **B-G)** Validating new ciliary candidates with additional stainings. Example images from stainings for the proteins indicated in hTERT-RPE1 cells (serum-starved) or RPTEC/TERT1 cells, using additional HPA or commercial antibodies targeting different immunogenic sequences (indicated in brackets where available). **H)** Example image from a staining of hTERT-RPE1 cells for CAMK1G and of hTERT-RPE1 cells overexpressing human CAMK1G fused to a C-terminal Halo tag. Both approaches show localization of CAMK1G at the Basal Body (BB) in hTERT-RPE1 cells.

#### Note S1: Patient Summary

The patient is a 2-year-old male, the second child of healthy, non-consanguineous parents from Sweden. His 5-year-old brother is healthy. There is a history of one early miscarriage before the patient was born. No family history of developmental anomalies, short stature, visual impairment, or congenital malformations was reported.

He was born at 38+1 weeks of gestation via vaginal delivery. Growth at birth was normal (Birth weight: 3550 g; length: 47 cm). At 3 months of age, he presented with severe early-onset vision impairment and progressive growth failure. At one year of age, height, weight, and head circumference had dropped from 0 to -4 SD. Eye examination revealed hypopigmented fundi with preserved central macula pigmentation, nystagmus, and normal-sized but slightly pale optic discs. An electroretinogram (ERG) showed findings consistent with retinal dystrophy of LCA type (Leber congenital amaurosis). Brain MRI findings revealed bilateral optic nerve hypoplasia (ONH), a smaller-than-expected optic chiasm, and a hypoplastic adenohypophysis (1 mm in height) with a shallow sella turcica. The hypothalamic regions appeared slightly atrophic. Additional structural anomalies include enlarged basal cisterns, a mega cisterna magna, wide cerebrospinal fluid spaces at the temporal base, and a small arachnoid cyst. A skeletal survey showed low bone density. He has no heart or kidney malformations, as confirmed by ultrasound.

By 1 year and 5 months, he was hospitalized due to severe hypoglycemia (0.6 mmol/L) leading to seizures. Repeated IGF-1 measures were low <15 ng/mL, TSH/fT4 within low-normal range (TSH 1.8; fT4 12 pmol/L), growth hormone (GH) was 0.7 ng/mL, cortisol levels were normal. He was diagnosed with ONH, growth hormone deficiency, and hypothyroidism. Hypoglycemia was considered secondary to GH deficiency. He started treatment with GH (Omnitrope) and levothyroxine, which improved his growth and well-being but not his vision.

Mild dysmorphic features include a high, prominent forehead, hypertelorism, prominent eyes with epicanthic folds, blue sclera, low nasal bridge, a short nose, and a thin upper lip. He also exhibited fair skin and hair, raising suspicion of albinism. Weight at 1 year and 3 months was 7.45 kg (-3.3 SD), height 65.5 cm (-4.23 SD), and head circumference 42.3 cm (-4.14 SD).

Clinical Genetic testing with trio whole genome analysis was negative, including analysis of all known genes associated with growth hormone deficiency, retinal dystrophies, or albinism. However, a research analysis detected a de novo two nucleotide deletion in CREB3 (NM_006368) (c.810_811del/p.Ser271*), causing a premature stop codon.

At 2 years and 6 months, the patient’s height has improved to -3 SD and no further hypoglycemia episodes have occurred. Vision remains severely impaired. Developmental progress includes standing, walking with support, and forming short sentences with several words. Hearing is normal.
